# *Drosophila* cells that express octopamine receptors can either inhibit or promote oviposition

**DOI:** 10.1101/2023.05.03.539296

**Authors:** Ethan W. Rohrbach, Elizabeth M. Knapp, Sonali A. Deshpande, David E. Krantz

**Affiliations:** Interdepartmental Program in Neuroscience, Brain Research Institute Gonda (Goldschmied) Neuroscience and Genetics Research Center, UCLA, Los Angeles, CA 90095; Department of Psychiatry and Biobehavioral Sciences, David Geffen School of Medicine, UCLA, Los Angeles, CA 90095

**Keywords:** octopamine, octopamine receptor, egg-laying, spermatheca, oviposition

## Abstract

Adrenergic signaling is known to play a critical role in regulating female reproductive processes in both mammals and insects. In *Drosophila*, the ortholog of noradrenaline, octopamine (Oa), is required for ovulation as well as several other female reproductive processes. Loss of function studies using mutant alleles of receptors, transporters, and biosynthetic enzymes for Oa have led to a model in which disruption of octopaminergic pathways reduces egg laying. However, neither the complete expression pattern in the reproductive tract nor the role of most octopamine receptors in oviposition is known. We show that all six known Oa receptors are expressed in peripheral neurons at multiple sites within in the female fly reproductive tract as well as in non-neuronal cells within the sperm storage organs. The complex pattern of Oa receptor expression in the reproductive tract suggests the potential for influencing multiple regulatory pathways, including those known to inhibit egg-laying in unmated flies. Indeed, activation of some neurons that express Oa receptors inhibits oviposition, and neurons that express different subtypes of Oa receptor can affect different stages of egg laying. Stimulation of some Oa receptor expressing neurons (OaRNs) also induces contractions in lateral oviduct muscle and activation of non-neuronal cells in the sperm storage organs by Oa generates OAMB-dependent intracellular calcium release. Our results are consistent with a model in which adrenergic pathways play a variety of complex roles in the fly reproductive tract that includes both the stimulation and inhibition of oviposition.

## INTRODUCTION

Oocyte development, fertilization, and ovulation are regulated by multiple steroid and peptide hormones as well as aminergic neuromodulators in both mammals and invertebrates. The regulation of some processes is surprisingly conserved, allowing the use of relative simple systems to explore the underlying mechanisms (White, Chen, and Wolfner 2021; Kamhi et al. 2017; E. M. Knapp et al. 2020; E. Knapp and Sun 2017). These include the adrenergic regulation of oocyte development and ovulation mediated by noradrenalin in mammals and its structural ortholog octopamine (Oa) in *Drosophila melanogaster* (Deady and Sun 2015; J. Kim et al. 2021; Hoshino and Niwa 2021; Yoshinari et al. 2020; Andreatta et al. 2018; J. Kim et al. 2021; L. Wang et al. 2022; J. Kim and You 2022; Čikoš et al. 2007; Schmidt et al. 1985; Kannisto, Owman, and Walles 1985; Kobayashi et al. 1983; Blum et al. 2004; Meiselman, Kingan, and Adams 2018).

In mammals, noradrenaline is known to directly modulate reproductive tract function (Lawrence and Burden 1980; M. T. Itoh et al. 2000; Masanori T. Itoh and Ishizuka 2005; Ricu et al. 2008) with proposed roles in ovarian steroid synthesis (Garrido et al. 2018; Adashi and Hseuh 1981) and muscle contractions (Virutamasen, Wright, and Wallach 1973; Kannisto, Owman, and Walles 1985; Kobayashi et al. 1983). Polycystic ovarian syndrome (PCOS) is a common anovulatory disease affecting millions of women in every region of the world (Wolf et al. 2018), and it has been shown to present with increased sympathetic signal release to the ovaries and elevated plasma noradrenaline levels (Lansdown and Rees 2012; Greiner et al. 2005). Determining conserved mechanisms by which adrenergic signaling regulates female fertility may aid in the development of novel therapeutic strategies for PCOS and other anovulatory conditions.

In *Drosophila* and other insects, the steps required for egg-laying (oviposition) include follicle development, follicle rupture, ovulation, passage through the oviducts and egg deposition (White, Chen, and Wolfner 2021; J. Sun and Spradling 2013; Häsemeyer et al. 2009; Feng et al. 2014; F. Wang et al. 2020; Deady and Sun 2015; Mattei et al. 2015; Avila et al. 2012; Orchard and Lange 1985; Lange 2009). Oa contributes to the regulation of most, if not all of these processes in *Drosophila* and other insect species (Kamhi et al. 2017; Andreatta et al. 2018; Orchard and Lange 1985; Lange 2009; Hana and Lange 2020; Wong and Lange 2014; Rezával et al. 2012; Yoshinari et al. 2020; White, Chen, and Wolfner 2021).

*Drosophila melanogaster* expresses six Oa receptors (Qi et al. 2017; Balfanz et al. 2005; Maqueira, Chatwin, and Evans 2005; Evans and Maqueira 2005; Farooqui 2007; El-Kholy et al. 2015). Two of these, *Oamb* and *Octβ2R*, have been established as critical for fertility and linked to several physiological processes involved in oviposition (H.-G. Lee et al. 2003; H.-G. Lee, Rohila, and Han 2009; Lim et al. 2014a; Deshpande et al. 2022; Deady and Sun 2015; Avila et al. 2012; Li et al. 2015). Expression of both receptors in the epithelium of the oviduct regulates egg laying (H.-G. Lee, Rohila, and Han 2009; Lim et al. 2014a). *Oamb* expression in ovarian follicle cells is required for follicle rupture (Deady and Sun 2015), and expression in parovarian organs as well as the seminal receptacle and spermathecae may play a role in sperm storage dynamics (Avila et al. 2012; 2010). We have recently shown that *Octβ2R* and *Oamb* are required for contraction and dilation of the lateral oviducts respectively (Deshpande et al. 2022). The effects of *Octβ2R* may be mediated via expression in either neurons (Deshpande et al. 2022) or muscle (Li et al. 2015). It is unclear whether *Oamb* or *Octβ2R* may regulate additional processes involved in oviposition, and both are widely expressed at multiple additional sites throughout the reproductive tract and CNS. The potential role for *Oamb* and *Octβ2R* expressing neurons in oviposition-linked circuits is particularly unclear, since most studies have focused on their function in non-neuronal tissues, such as the epithelium and follicle cells (H.-G. Lee, Rohila, and Han 2009; Lim et al. 2014a; H.-G. Lee et al. 2003; Li et al. 2015; Deady and Sun 2015; White, Chen, and Wolfner 2021). With regard to the other four Oa receptors expressed in *Drosophila* -- *Octα2R*, *Octβ1R*, *Octβ3R* and *Oct-TyrR*—it is not known whether they play any role in oviposition, or in which type(s) of cells in the reproductive tract they may be expressed (El-Kholy et al. 2015). Here and in a previous report, we have used a panel of molecular tools for high-fidelity expression of each receptor to determine where they are expressed in the reproductive tract and how cells expressing each one may contribute to oviposition (Deshpande et al. 2022).

Although Oa regulates multiple processes within the reproductive tract, the most commonly reported phenotype for mutations that reduce octopaminergic signaling is a decrease in the release of eggs from the ovary (ovulation) with a resultant decrease in egg laying (White, Chen, and Wolfner 2021; Pang et al. 2022). This phenotype has been observed with loss of function mutants or RNAi for the receptors *Oamb* (H.-G. Lee et al. 2003; H.-G. Lee, Rohila, and Han 2009; Deady and Sun 2015) and *Octβ2R* (Lim et al. 2014a; Li et al. 2015), an enzyme required for Oa synthesis, tyrosine β- hydroxylase (TβH) (Monastirioti, Charles E. Linn, and White 1996; Cole et al. 2005; Monastirioti 2003) and the transporter responsible for its storage and release from secretory vesicles (Simon et al. 2009). Oa also has the potential to influence fertility via actions in the oviducts or uterus (Deshpande et al. 2022; Lim et al. 2014b; Li et al. 2015; H.-G. Lee, Rohila, and Han 2009; H.-G. Lee et al. 2003); however, effects that occur in the uterus or other sites downstream of follicle rupture can be epistatically occluded by retention in the ovaries.

Both gain of function and loss of function mutations can be useful for genetic analyses, e.g., for epistatic experiments to determine the order of genetic or biochemical events. With perhaps one notable exception (Hoff et al. 2011), gain of function transgenes for *Drosophila* Oa receptors are not available. Moreover, over-expressing a receptor may not increase the activity of post-synaptic neurons if the presynaptic input is not amplified. Oa receptors are Gα_s,_ and Gα_q_ coupled GPCR’s, and directly activating neurons in which the Oa receptors are expressed represents an alternative approach to probe their potential functions. The molecular tools available in *Drosophila* are well suited to drive expression of probes that activate or inhibit specific subsets of neurons, including the dTrpA1 Ca^2+^ channel and the Kir2.1 K^+^ channel. These and other probes can be easily combined with high fidelity drivers such as “MiMICS”, in which a Gal4 or LexA transcription factor is embedded within the endogenous receptor gene to precisely replicate its temporal and spatial patterns of expression (Diao et al. 2015; Venken et al. 2011; McKinney et al. 2020; Deshpande et al. 2022; P.-T. Lee et al. 2018).

We have used a panel of six MiMIC lines to map the expression in the reproductive tract and the potential function(s) in oviposition of the six known *Drosophila* Oa receptors (McKinney et al. 2020; Deshpande et al. 2022). We demonstrate co-expression of Oa receptor subtypes with the mechanosensitive marker *ppk1.0* and the acetylcholine marker *ChAT* in uterine cells previously shown to regulate oviposition (Yoshinari et al. 2020; Yang et al. 2009; Gou et al. 2014; F. Wang et al. 2020; Deshpande et al. 2022; Rezával et al. 2014; 2012). Activation of neurons that express at least three of the receptors stimulates oviduct contraction, and we show that *Oamb* regulates the acute response of cells in the sperm storage organs to Oa. Behavioral assays indicate that activation or inhibition of neurons that express each receptor impacts oviposition in a different way, but in contrast to most previous studies in which Oa has been suggested to promote oviposition (Deady and Sun 2015; Cole et al. 2005; Li et al. 2015; H.-G. Lee, Rohila, and Han 2009; Lim et al. 2014a; Simon et al. 2009; Monastirioti, Charles E. Linn, and White 1996; J. Kim et al. 2021; Hoshino and Niwa 2021; White, Chen, and Wolfner 2021), our data suggest a complementary role for octopamine receptor-expressing neurons (OaRNs) in the inhibition of egg-laying. This includes a potentially novel role for *Oamb*-expressing neurons to promote egg retention in the uterus.

## MATERIALS AND METHODS

### Experimental model and subject details

Flies were raised in mixed sex vials on cornmeal/sucrose/yeast/sucrose/dextrose/agar medium at 25°C and 50–70% humidity under a 12:12 light: dark cycle unless otherwise noted. All experiments used mated or virgin female flies 4–6 days post eclosion. All fly lines used in this study are listed in Supplementary Table S1.

### Fly Husbandry and stocks

Publicly available fly lines with noted identifiers were obtained from the BDSC (Listed in Supplementary Table 1). We thank the following people for generously supplying the following additional lines: Kyung-An Han (University of Texas, El Paso) for *OAMB-RS-GAL4*; Bing Ye (University of Michigan) for *ppk1.0-LexA* and *ppk-Gal4* (Gou et al., 2014), Soo Hong Min (Harvard) for *elav-Gal80* (Yang et al. 2009), Robert Kittel (University of Würzberg) for *UAS-ChR2-XXM* (Dawydow et al. 2014; Scholz et al. 2017), and Mark Frye (UCLA) for *UAS-Kir2.1* and *tub-Gal80^ts^*.

### Immunofluorescence staining

To visualize the expression patterns of the Oa receptor genes, complete reproductive systems from flies harboring *UAS-mCD8-GFP* and either Oa receptor *MiMIC-T2A-Gal4* lines or *ChAT-Gal4* were dissected in phosphate buffered saline (PBS) at 25 °C with each innervating VNC and MAN connection left intact. The tissue was fixed in 4% paraformaldehyde (PFA) for 15 min and washed in PBS containing 0.3% (vol/vol) Triton X-100 (PBT) for 30 minutes. Following fixation, preparations were washed for 1 hour in blocking buffer containing 5% (vol/vol) normal goat serum (NGS) (Cat# G9023, Sigma-Aldrich) in PBT then incubated for 24–48 hr at 4 °C in the primary antibody mouse anti-GFP (1:500 in blocking buffer, Ref# A11120, Invitrogen). The sample was washed in PBT for 3 hr at 25 °C then incubated in the secondary antibody AF488-conjugated goat-anti-mouse (1:500 in blocking buffer, Ref# A21202, Invitrogen) at 4 °C for 24 hr. After washing in PBT for 1 hr at 25 °C, the preparations were optically cleared in 25% glycerol for 1 hr at 25 °C and then mounted on Superfrost slides (Cat# 12-550-143, Fisherbrand) with bridged Glass Cover Slips #0 (Cat# 72198-10, Electron Microscopy Sciences) in Fluoromount-G mounting media (Cat# 0100-01, SouthernBiotech). For co-staining of muscle cells, preparations were dissected and processed as described above, except AF555-conjugated Phalloidin stain (1:500, Ref# A34055, Invitrogen) was included in the secondary antibody solution. For co-labeling of *ppk1.0-LexA*, preparations were dissected and processed as described above, except rabbit anti-dsRed (1:500, Cat# 632496, Takara Bio) was added to the primary antibody solution and AF568-conjugated goat-anti-rabbit (1:500, Ref#A31572, Invitrogen) was added to the secondary antibody solution. For co-staining of neuronal cells, preparations were dissected and processed as described above, except Rhodamine (TRITC) conjugated rabbit anti-horseradish peroxidase (HRP) (1:500, Code# 323-025-021, Jackson Immunoresearch) was included in the secondary antibody solution.

### Optogenetic stimulation of lateral oviduct contractions

For optogenetic experiments, mated female flies were raised in standard food containing 80 μM all-trans retinal from 1 day post eclosion until tested (5-7 days). Optogenetic stimulations of OaRNs were performed on “Abdominal Fillet” preparations as previously described (Deshpande et al. 2022). In brief the abdomen was separated from rest of the fly body using microscissors, pinned to a Sylgard dish and the sternal plates removed to expose the reproductive organs. Stimulation was performed using a Lambda DG-4 light source (Sutter), Chroma filter set 41001 and the light path of an AxioExaminer Z1 microscope to illuminate the entire field of view at 1mW/mm^2^ power. The preparation was also illuminated from the side using an external LED to facilitate the visualization of contractions. Contractions of the LO were defined by decrease in the distance between ovaries and a characteristic contraction of the oviduct tissue as described (Deshpande et al. 2022) and counted in videorecordings using a CCD camera (Andor iXon 897, Oxford Instruments, Oxfordshire, England) at a capture rate of 12 frames/sec and using Andor IQ2 software. Flies harboring one copy of *Octβ1R*, *Octβ3R*, or *Oct-TyrR MiMIC-T2A-Gal4* and one copy of *UAS-ChR2-XXM::tdTomato* were compared to control flies with one copy of the *MiMIC-T2A-Gal4* that had been crossed with *Canton-S* (CS) flies.

### Live imaging of sperm storage organs

To determine Oa’s effects on the accessory gland cellular activity, we prepared fly reproductive tracts for live imaging using the previously described “Isolated Preparation” method (Deshpande et al. 2022). The abdominal cuticle, the gut and fat bodies was removed to generate an intact, “isolated” reproductive tract. The anterior tip of the ovaries and the distal end of the uterus were pinned to a Sylgard substrate with insect pins. Instead of focusing on oviduct regions as in (Deshpande et al. 2022), we recorded images of the accessory glands including the seminal receptacle and spermathecae. Live imaging of RCaMP1b was performed using a 555 nm LED light source (Thorlabs), the standard Chroma filter set 41007a, a Zeiss Achroplan water immersion 10x objective on a Zeiss Axio Examiner Z1 microscope with a CCD camera (Andor iXon 897, Oxford Instruments, Oxfordshire, England) at a capture rate of 12 frames/sec using Andor IQ2 software. Images were analyzed using Fiji/ImageJ software (Schindelin et al. 2012). For all Regions of Interest (ROIs), an off-target area of equal size was selected as background. Changes in fluorescence are reported as the background-subtracted difference in the change in fluorescence divided by baseline (ΔF/F = [(F peak - F baseline)/F baseline], where F baseline = average RCaMP signal during the 1 min before Oa bath application). In all experiments, 1 min of baseline activity was recorded before Oa was bath applied to the preparation at the indicated concentration. Recording was conducted for 4 min following Oa addition. All flies tested were 4 days post eclosion and carried either one copy of *24b-Gal4* or *40B09-Gal4* and one copy of *UAS-RCaMP1b*. RNAi experiments used flies that additionally included one copy each of *UAS-dicer2* and *UAS-OAMB-RNAi*. Mated flies were cohoused with *CS* males after sorting 1 day post eclosion. Virgin flies were sorted and housed without males.

### Fertility assays

Egg laying assays were conducted as previously described (E. Knapp and Sun 2017). In brief, virgin flies harboring the indicated *Gal4* alleles and either *UAS-TrpA1*, *UAS-Shibire^ts^*, or *UAS-Kir2.1* and *TubGal80^ts^* were collected and housed at 22 °C. At 5 days post eclosion, virgins were mated with *CS* males and either kept at 22 °C for control experiments or shifted to 29 °C to facilitate hyperactivation or suppression of neural activity via transgene expression. The number of eggs laid on molasses plates were then counted every day for 2 days and averaged. Egg laying times were calculated by dividing 1440 min by the total number of eggs laid per female per day. Following the 2 day collection of egg laying data, the ovaries from each fly were dissected for quantification of mature follicles as previously described (E. Knapp and Sun 2017): ovaries were fixed in 4% PFA for 15 min, stained with DAPI (1:1000, Prod#62248, Thermo Scientific), and mounted on Superfrost slides with 24x30-1 Cover Glass (Fisherbrand). Mature follicles in each set of ovaries were quantified via fluorescent imaging using a Zeiss LSM 880 confocal microscope. To partition egg laying times into ovulation time, oviduct time, and the uterus time, partitioning ratios were determined by dissecting reproductive tracts after separate 6 hr mating experiments and determining the percentage of females with eggs in the oviduct or uterus as previously described (E. Knapp and Sun 2017).

## RESULTS

### Multiple subtypes of octopamine receptors are expressed by central and peripheral neurons that innervate the reproductive tract

We visualized the expression of each one of the six *Drosophila* Oa receptors using a panel of *MiMIC-T2A-Gal4* lines (McKinney et al. 2020; Diao et al. 2015; Venken et al. 2011). Consistent with previous reports (H.-G. Lee et al. 2003), we detect expression of *Oamb* in follicle cells that surround the mature oocyte (Fig. 1, Bi, filled arrowhead) and expression of both *Oamb* and *Octβ2R* in epithelial cells that line the lumen of the oviducts (Fig. 1, Bi, Bv, open arrowheads). We did not detect the expression of Oa receptors other than *Oamb* in follicle cells (Fig. 1 and data not shown). Functional data also support a role for *Oamb* in somatic escort cells in regulating the germline stem cell lineage, exclusive of its role in ovulation and follicle cell rupture (Yoshinari et al. 2020). We did not assess the potential expression of other Oa receptors in intra-ovariole cells, and this will require further study.

**Figure 1.**
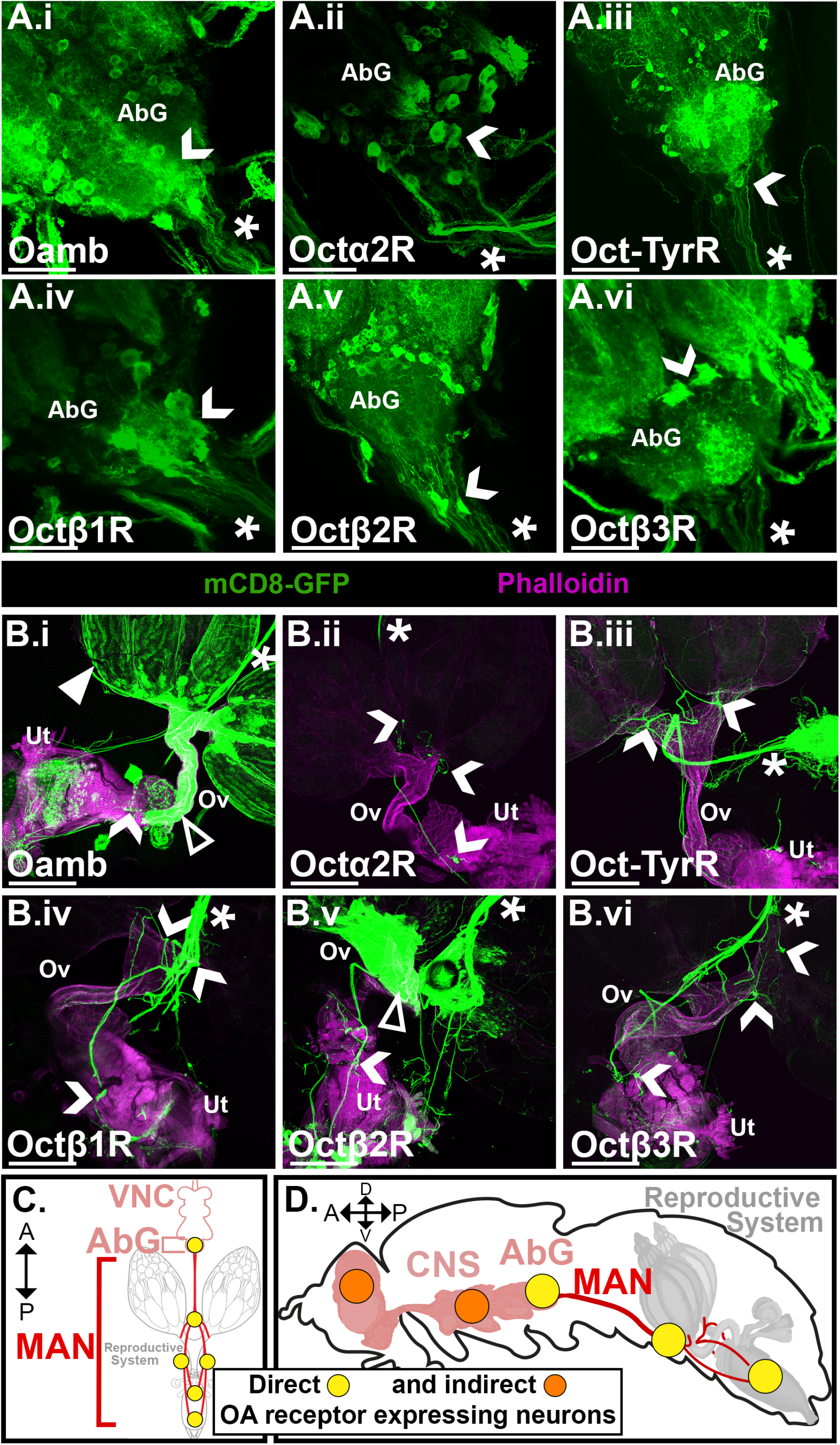
Octopamine receptor expression in cells associated with the reproductive system. A panel of Oa receptor drivers (*MiMIC-T2A-Gal4* lines) was used to drive *UAS-mCD8-GFP*. For each genotype, the Abdominal ganglion (AbG) (**A**) and the reproductive system (**B,** Ov=oviduct, Ut=uterus) were dissected then labeled with anti-GFP ALEXAfluor-488 (green) and Phalloidin-555 (magenta) to visualize Oa receptor expressing cells and muscles respectively. Labeled nerves in the medial abdominal nerve (MAN, stars) project from neurons in the CNS to reproductive system (**A,** chevrons) and from neurons in the reproductive system back to the CNS (**B,** chevrons). *Oamb* is also expressed in follicle cells (filled arrowhead). Both *Oamb* and *Octβ2R* are expressed in epithelial cells (open arrowheads). **C.** Ventral-view cartoon depicting the approximate locations of neuronal clusters expressing Oa receptors directly associated with the reproductive system. **D.** Sagittal-view cartoon showing the approximate locations of groups of Oa receptor expressing neurons in the full CNS and reproductive system. Orange circles indicate clusters of cells known to express Oa receptors that are not directly associated with the reproductive system but could potentially influence oviposition. Scale bars = 20μm (**A**) and 200μm (**B**)

We and others have previously shown that *Octβ2R* is expressed in multiple neurons within the reproductive system as well as neurons in the CNS that descend from the abdominal ganglion (AbG) (Deshpande et al. 2022; McKinney et al. 2020). We find that *Octα2R*, *Octβ1R*, *Octβ3R*, and *Oct-TyrR* expressing cells also extend processes throughout the reproductive system (Fig. 1, A, B, chevrons, C, D, yellow circles, see also Supplemental Fig. 1). Boutons in the reproductive tract that express different types of Oa receptors appear to have distinct morphologies (Sup. Fig. 1H). We also show quantitative differences in the innervation of specific organs within the reproductive tract by each OaRN subtype (Sup. Fig. 1, G). Based on labeling with an antibody to a neuron specific epitope (“anti-HRP”/nervana), most of these processes appear to be neuronal (Supplemental Fig. 2).

As others have reported, we detect neuronal cell bodies that express *Oamb, Octα2R, Octβ1R, Octβ2R, Octβ3R and Oct-TyrR* in the Abdominal Ganglion (AbG) (Fig. 1, A, chevrons) (McKinney et al. 2020). We further confirm that projections from these neurons extend into the median abdominal nerve (MAN) that connects the AbG to the reproductive system (Fig. 1, A, B, stars).

We also observe multiple peripheral neurons that express *Oamb, Octα2R, Octβ1R, Octβ2R, Octβ3R and Oct-TyrR* and are embedded within the MAN and/or the reproductive tract itself (Fig. 1, B, chevrons). Cells embedded within uterine muscle layers appear to project ascending processes through the MAN into the CNS, whereas cells embedded within the MAN are difficult to distinguish as either afferent or efferent (Fig. 1, chevrons). Single cell labeling methods will be necessary to definitively determine the projection patterns and connectivity of each neuron. In contrast to *Oamb* and *Octβ2R,* we do not detect expression of *Octα2R, Octβ1R, Octβ3R or Oct-TyrR* in any of the epithelial cells that line the lumen of the oviducts (Fig. 1 and data not shown).

### Cholinergic ppk1.0(+) neurons express multiple subtypes of octopamine receptors

Sex peptide (SP) released by males during copulation regulates the post-mating response of females via disinhibition of oviposition-promoting signals (P. S. Chen et al. 1988; Aigaki et al. 1991; Soller, Bownes, and Kubli 1997; Chapman et al. 2003; Häsemeyer et al. 2009; Yang et al. 2009; Avila et al. 2010; Yoshinari et al. 2020). The set of neurons that expresses the SP receptor gene (*SPR*) is also known to include peripheral neurons that express a form of acid-sensing sodium channel ppk, ppk1. The “*ppk1.0*” driver includes a fragment of the *ppk1* regulatory DNA and is expressed in a subset of *ppk1*(+) neurons in the reproductive-tract (Gou et al. 2014; Häsemeyer et al. 2009; Greenspan 1980). Importantly, the “*ppk1.0*” labeled subset of neurons have been shown to be involved in the post-mating response (Gou et al. 2014; Häsemeyer et al. 2009; Greenspan 1980). We previously demonstrated co-expression of *ppk1.0-LexA* with the *MiMIC-T2A-Gal4* drivers representing *Octβ2R* and *Oamb* (Deshpande et al. 2022). Here we again used the *ppk1.0-LexA* driver to determine if other Oa receptors are expressed in *ppk1.0*(+) cells within the reproductive tract, focusing on a specific cluster of three stereotypically localized *ppk1.0*(+) neurons in the anterior uterus. Since some *ppk1*(+) cells are known to be a subset of cholinergic, *SPR*(+) neurons, we also compared *ppk1.0* expression to *ChAT-Gal4* expression to better contextualize individual neuron identities among peripheral clusters (Fig. 2 A) (Yoshinari et al. 2020; Grueber et al. 2007; 2003).

**Figure 2.**
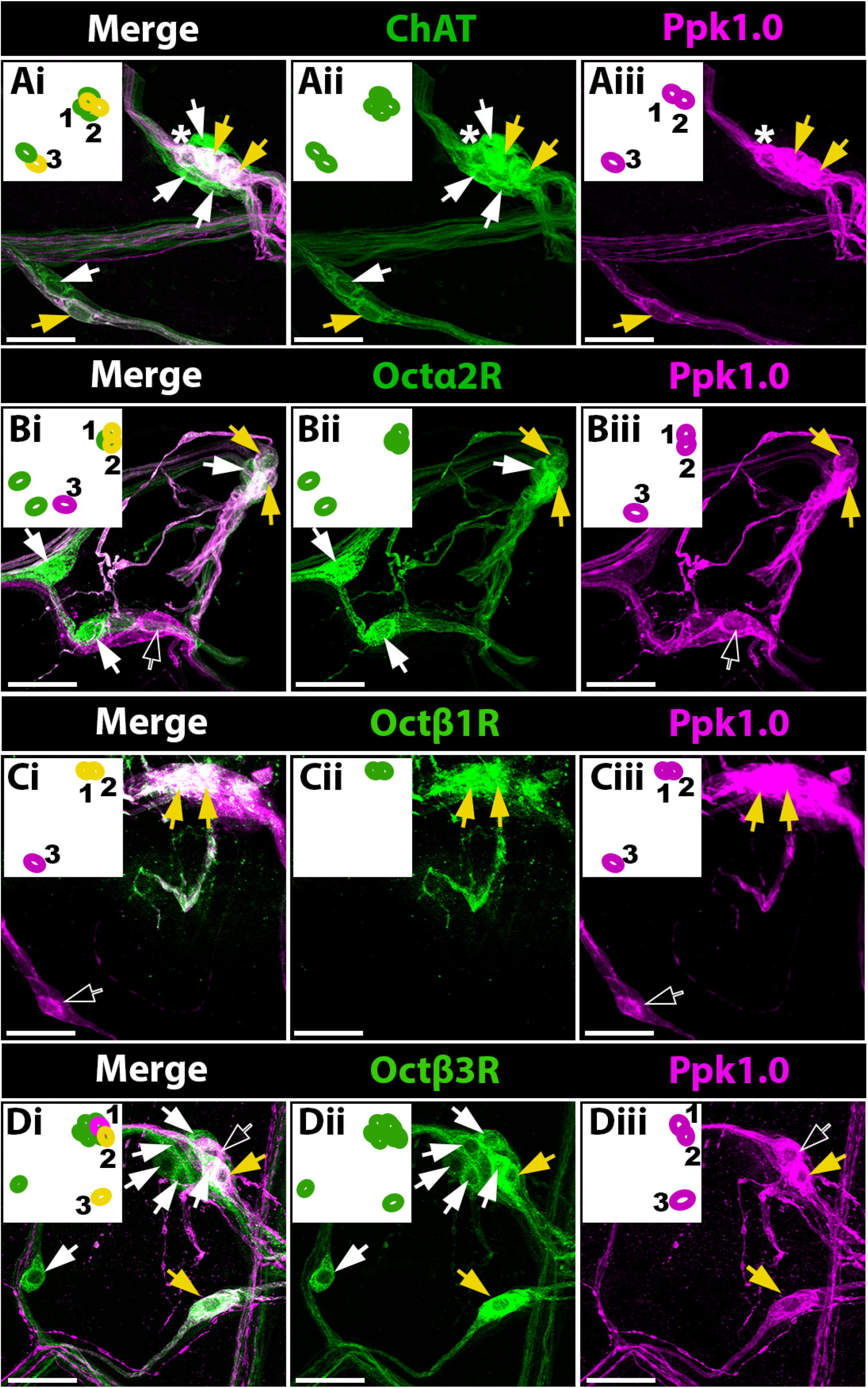
Peripheral *ppk1.0*(+) neurons express octopamine receptors and are cholinergic. Octopamine receptor MiMIC Gal4’s or *ChAT-Gal4* were used to drive *UAS-mCD8::GFP* and *Ppk1.0-LexA* was used to drive *LexAop-CD2-RFP*. *ChAT* (**A**) *Octα2R* (**B**), *Octβ1R* (**C**), *Octβ3R* (**D**) expression was assessed among three stereotyped *ppk1.0*(+) cell bodies embedded within the anterior uterus. Cells that co-express of *ppk1.0-LexA* with each of the *Gal4* drivers are indicated (yellow arrows). Also indicated are [*ppk1.0(+)*, *Gal4(-)*] cells (black arrows), [*ppk1.0(-)*, *Gal4*(+)] cells (white arrows), and a site where *ppk1.0(+)* and *Chat(+)* processes overlap (A, asterisk). Images are maximum signal projections through 20μm. Insets show cell body locations expressing each marker with the three stereotyped *ppk1.0*(+) cell bodies numbered 1-3 and yellow ovals in the “Merge” insets indicating co-expression. Scale bars = 10 μm

Among three stereotypically localized *ppk1.0(+)* neurons embedded within the muscle cells that envelop the anterior uterus (Fig. 2Aiii - Diii), we confirm that all three also express *ChAT-Gal4 (*Fig. 2A, yellow arrows). We observe *ChAT* expression in four adjacent cells that do not express *ppk1.0-LexA* (Fig. 2A, white arrows). It is possible that these cells express *ppk1* (and *SPR*) but are not included in the more restricted subset of cells labeled by the *ppk1.0* driver. Additionally, though it may appear as if there is a third *ppk1.0(+)* neuron in these images (Fig. 2 A, white asterisk), single-slice image analyses as well as analyses of additional preparations show that there are indeed only three ppk1.0(+) cell bodies in this region (data not shown).

Of these three *ppk1.0(+)* neurons, two consistently localize together (Fig. 2, inset, cells “1” and “2”) while a third is consistently seen as separated from the other two by approximately 20 μm (Fig. 2, cell “3”). The two *ppk1.0(+)* cells that exist side-by-side (“1” and “2”) are *Octα2R*(+) and *Octβ1R(+)* (Fig. 2B, C, yellow arrows). The isolated *ppk1.0(+)* neuron (“3”) does not detectably express *Octα2R* (2B, black arrow) or *Octβ1R(+)* (2C, black arrow). In contrast to *Octα2R* and *Octβ1R(+), Octβ3R is* clearly expressed in the isolated *ppk1.0(+)* neuron (Fig. 2D, lower right, yellow arrow) and at least one of the side-by-side *ppk1.0(+)* neurons (“1” or “2”) (Fig. 2D, upper right, yellow arrow). These data suggest that activation of *ppk1.0(+)* neurons that express Oa receptors could inhibit egg laying similarly to the activation of *SPR(+)* neurons (P. S. Chen et al. 1988; Aigaki et al. 1991; Soller, Bownes, and Kubli 1997; Chapman et al. 2003; Häsemeyer et al. 2009; Yang et al. 2009; Avila et al. 2010; Yoshinari et al. 2020). Furthermore, our observation that specific *ppk1.0(+)* neurons express different sets of Oa receptor subtypes suggests the possibility that Oa could differentially regulate their activity.

### *Oamb* is expressed in non-neuronal secretory gland and uterine cells

In a previous report, we focused on *Oamb* expression in relatively anterior regions within the reproductive tract and the role of *Oamb* in lateral oviduct contractility (Deshpande et al. 2022). Oamb also regulates the function of more posterior regions of the reproductive tract, including the function of the spermathecae, one of two sperm storage organs in the fly (Avila et al. 2012). The second sperm storage organ is the seminal receptacle, a compact, muscular tube attached to the uterus, also regulated by Oa (Avila et al. 2012). The parovarian glands are bilaterally symmetric structures posterior to the spermatheca and thought to also play a role in hormone secretion, however, their function remains poorly understood (Ito and Tomioka 2016; Wilson et al. 2017; Peng et al. 2005; Avila et al. 2012; Findlay et al. 2014; Ram and Wolfner 2009; Claudia Fricke et al. 2013; C. Fricke et al. 2009). Similar to a previous study (H.-G. Lee et al. 2003), the *Oamb* MiMIC line shows expression in the parovarian glands (Fig. 3, C). However, we also find robust expression of *Oamb-T2A-Gal4* at several sites within the sperm storage organs including the spermathecal secretory cells (Fig. 3, B), cells in the seminal receptacle that appear to represent an epithelial layer luminal to the muscle (Fig. 3, A), and neuronal processes that innervate each organ (Fig. 3, A-C, stars). We also detect a cluster of small cells expressing *Oamb* embedded between the muscle cells of the posterior uterus (Fig. 3, D). The cells in the posterior uterus extend processes from their somata and thus appear morphologically neuronal; however, a subset of at least three such *Oamb*(+) cells is not detectably labeled with the commonly used neuronal marker anti-HRP/nervana (B. Sun and Salvaterra 1995), and could potentially represent another cell type (Fig. 3, E, F, G).

**Figure 3.**
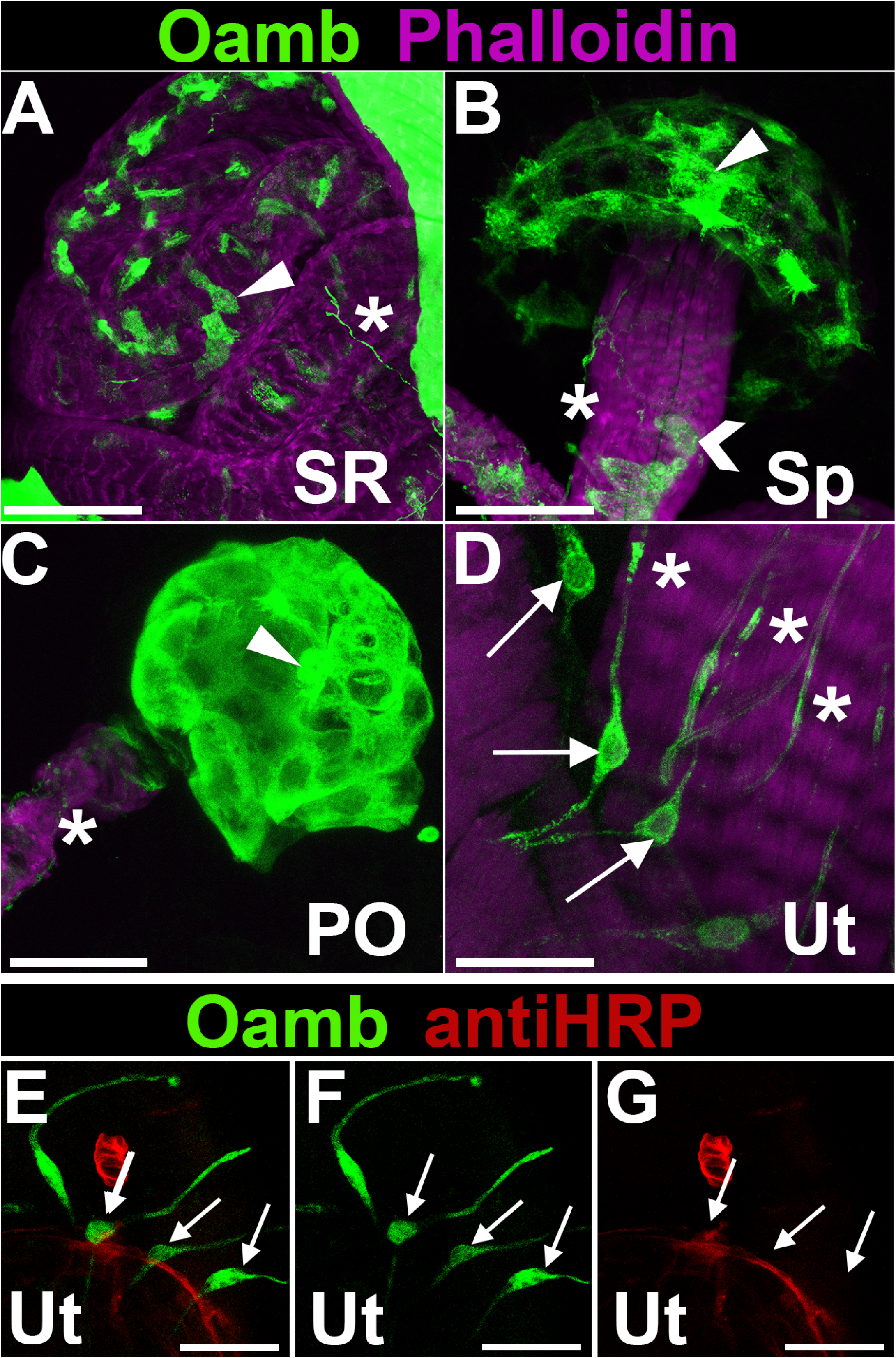
*Oamb* is expressed in the seminal receptacle, spermathecae and posterior uterus. *Oamb-T2A-Gal4* was used to drive *UAS-mCD8-GFP*. For each genotype, the reproductive system was dissected then labeled with anti-GFP ALEXAfluor-488 (green) and Phalloidin-555 (magenta) to visualize *Oamb* expressing cells and muscles respectively. We find *Oamb* expression in non-muscle cells in the seminal receptacle (**A,** SR), spermathecae (**B,** Sp), paraovia (**C,** PO), and posterior uterus (**D,** Ut). These include thin processes that may be neuronal (**A-D**, asterisks), cells that appear to be luminal to the muscle in the SR (**A**, arrowhead), secretory cells in the bulb of the Sp (**B**, arrowhead), epithelium like cells in the stalk of the Sp (**B**, chevron), cells in the PO of unknown function (**C**, arrowhead) and cell bodies in the uterus associated with thin possibly neuronal processes (arrows). Despite their neuronal-like morphology, the posterior uterus cells and processes do not label with anti-HRP (**E-G**). Scale bars = 10μm

### Optogenetic stimulation of OaRNs drives lateral oviduct muscle contractions

We have previously shown that optogenetic stimulation of neurons that express *Octβ2R* induces lateral oviduct contractions (Deshpande et al. 2022). To determine whether the other OaRNs might also promote oviduct contractility, we expressed the channelrhodopsin variant *ChR2-XXM* using each of the Oa receptor MiMIC Gal4 lines (Dawydow et al. 2014; Scholz et al. 2017). We employed a previously described assay in which optogenetic stimulation of octopaminergic neurons was paired with quantitation of lateral oviduct contractions (Fig. 4) (Deshpande et al. 2022). We used a preparation in which the AbG had been removed and the reproductive tract remained in situ within the abdomen (Fig. 4, A; an “Abdominal Fillet” see (Deshpande et al. 2022) and Methods). In addition to *Octβ2R*(+) neurons, we find that optogenetic activation of *Octβ1R*(+) and *Oct-TyrR*(+) neurons can reliably induce lateral oviduct contractions. Stimulation of two of five preparations using the *Octβ3R* MiMIC driver was followed by lateral oviduct contractions (Fig. 4, B). Importantly, although the oviduct epithelium expresses both *Octβ2R* and *Oamb*, we have previously shown that optogenetic stimulation of epithelial cells has no detectable effects on oviduct muscle contractions (Deshpande et al. 2022). In addition, we do not detect expression of any Oa receptors in the muscles of the reproductive tract (Fig. 1 and (Deshpande et al. 2022)). We therefore conclude that cells expressing *Octβ1R, Octβ3R and Oct-TyrR* that respond to optogenetic stimulation are likely to be neurons, although it is not possible to rule out another, novel type of electrically excitable cell.

**Figure 4.**
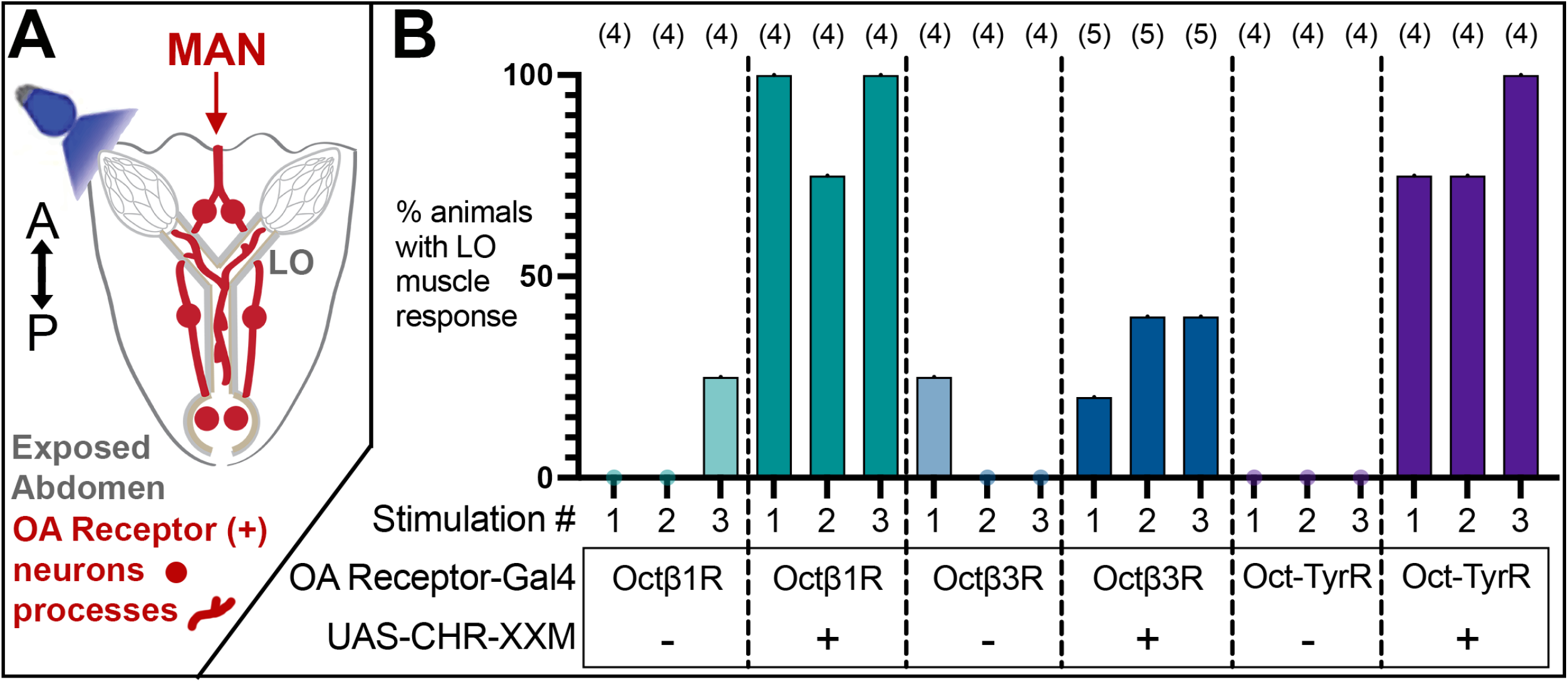
Optogenetic stimulation of octopamine receptor expressing neurons drives lateral oviduct (LO) contractions. **A.** Fly abdomens were dissected to expose the reproductive organs, and neurons with cell bodies or processes in the periphery were optogenetically stimulated **B.** The probability of a LO contraction response to each of 3 successive stimulations per fly was then quantified. Stimulation of *Octβ1R-T2A-Gal4* and *Oct-TyrR-T2A-Gal4* expressing cells reliably induces LO contractions. Stimulation of *Octβ3R-T2A-Gal4* expressing cells produces LO contractions less consistently. In control flies (without *UAS-ChR2-XXM*) we detected two instances of presumably, spontaneous contractions during periods of light exposure.

### Bath applied octopamine elicits Oamb-dependent global calcium changes in sperm storage organs

In addition to optogenetic stimulation of *Tdc2*(+) and OaRNs, bath applied Oa induces muscle contractions and calcium transients within the muscle cells of both the ovaries and the lateral oviducts (Middleton et al. 2006; Deshpande et al. 2022; Meiselman, Kingan, and Adams 2018). It has been hypothesized that epithelial expression of *Oamb* or *Octβ2R* in the oviduct may facilitate Oa-dependent regulation of muscle behavior (Lim et al. 2014). Muscle cells that can be labeled with the *24B-Gal4* driver (Martínez-Azorín et al. 2013) also surround the lumen of the seminal receptacle, where we find expression of *Oamb* in an epithelial-like layer but not in muscle (see Figure 3A). To determine if Oa could affect the muscles of the seminal receptacle, we used *24B-Gal4* to express *UAS-RCaMP1b* and quantitated changes in fluorescence (ΔF/F) at this site as a measure of cytosolic calcium at baseline and in response to bath applied Oa. We observed frequent spontaneous calcium transients at baseline in the muscle cells of the seminal receptacle (Supplementary Video 1). We also observed an increase in global calcium levels in the seminal receptacle following application of Oa (Fig. 5, A, B and Supplementary Video 1). Interestingly, calcium transients in the seminal receptacle appeared as waves, both at baseline and in the presence of Oa (Supplementary Video 1). In contrast to the increase in the intensity of the *RCaMP* signal, the rate of the calcium waves was not detectably altered by Oa bath application (data not shown). In similar experiments, we were unable to detect any changes in fluorescence or contractions within the muscle layers of the uterus (data not shown), although we cannot rule out a subtle effect on muscle activity at this site.

**Figure 5.**
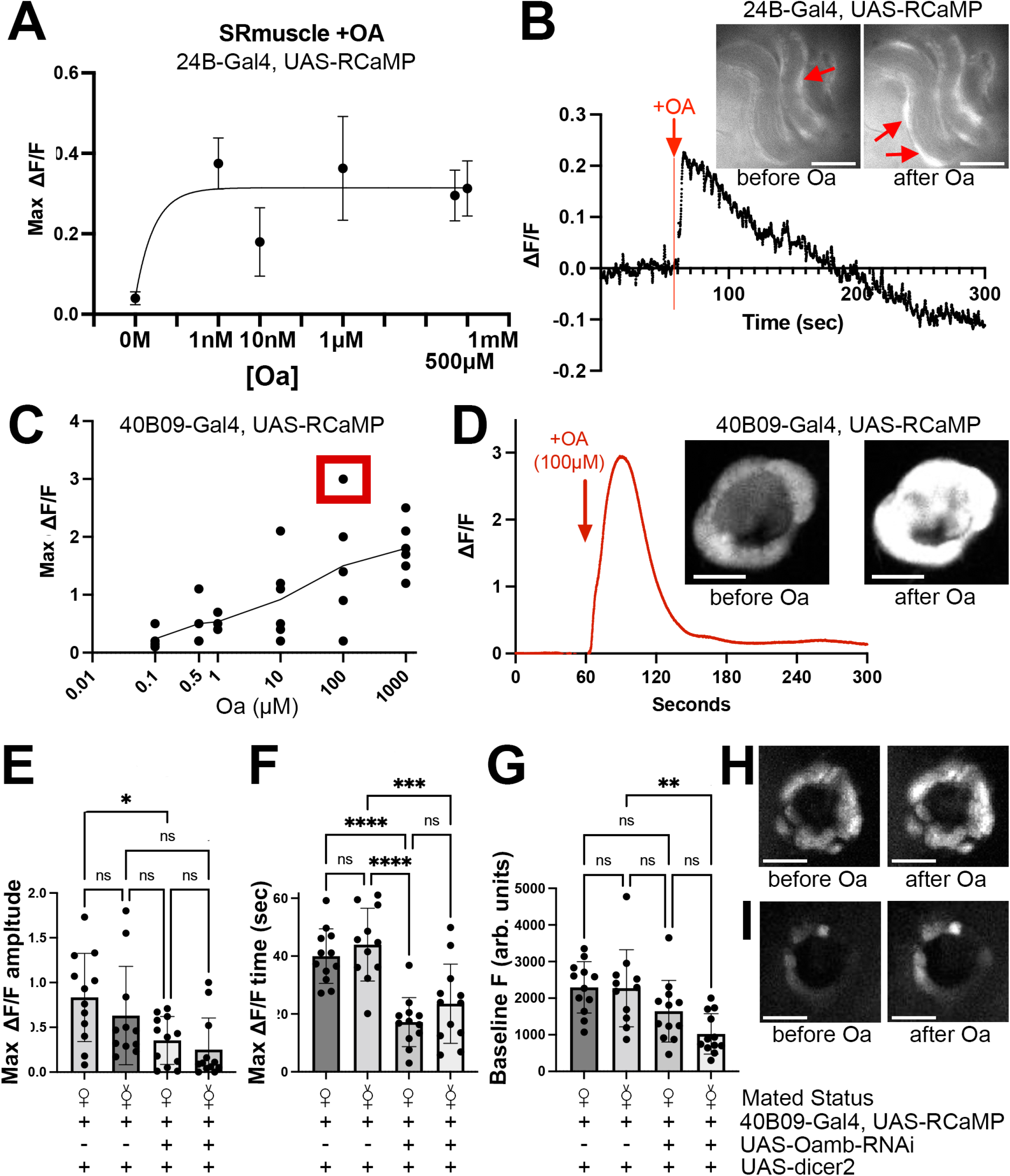
*Oamb* mediates sperm storage gland responses to octopamine. **A-B.** *UAS-RCaMP1b* expression was driven by the muscle driver *24B-Gal4*, and Oa was applied to reproductive systems dissected in HL3.1. **A.** Seminal receptacle muscle cells increase free intracellular calcium in response to Oa doses ranging from 1nM to 1mM. **B.** Sample trace and images from a recording in which 1μM Oa was delivered. **C-G.** *UAS-RCaMP1b* expression was driven by the spermatheca secretory cell driver *40B09-Gal4*, and Oa was again applied to reproductive systems dissected in HL3.1. **C.** The secretory cells of the spermathecae robustly increase free intracellular calcium levels in response to Oa doses ranging from 100nM to 1mM. **D.** Example trace (**C**, red box) including before and after images. Knocking down *Oamb* in the spermathecae secretory cells significantly reduces the maximal RCaMP response to Oa in mated flies (**E**) reduces the reduces the time to max response in both virgin and mated flies (**F**) and reduces the baseline RCaMP signal in virgin flies (**G,** arbitrary units). The distribution of the RCaMP signal at baseline (**H, I**, “before Oa”) and after adding Oa (**H, I**, “after Oa”) appeared to be less evenly distributed in both mated (**H**) and virgin (**I**) *Oamb* knockdown flies compared to those expressing WT levels of *Oamb* (**D**). Scale bars = 50μm

### Oamb elicits calcium transients in secretory cells within the spermatheca

The secretory cells in the spermathecae gland have been suggested to be regulated by Oamb (Avila et al. 2012) and previously shown to be involved in sperm storage (Schnakenberg, Matias, and Siegal 2011; Allen and Spradling 2008; Filosi and Perotti 1975). As shown above (Fig. 3, B), we find that *Oamb* is expressed in the spermathecae in both neuronal and non-neuronal cell types, including the secretory cells. We did not detect the expression of any Oa receptors other than *Oamb* in the non-neuronal cells within these organs (Fig. 1, B, and data not shown) suggesting that Oamb could be solely responsible for their response to Oa.

To test the acute effects of Oa on the secretory cells of the spermathecae we performed a series of calcium imaging experiments using RCaMP1b. We expressed *UAS-RCaMP1b* using a Gal4 driver (*40B09-Gal4*0 that is expressed specifically in the secretory cells of the spermathecae as demonstrated by the baseline RCaMP signal (Fig. 5, D). We then bath applied Oa (or vehicle) to “Isolated Preparations” of the reproductive tract that had been dissected out of the abdomen as described (Deshpande et al. 2022) (see Methods). We observe a dose-dependent increase in cytosolic calcium within *40B09-Gal4* expressing cells in response to Oa at concentrations as low as 100nM (Fig. 5, C, D).

To directly assess the contribution of *Oamb* to calcium transients in spermathecal cells, we used a previously tested RNAi transgene directed against *Oamb* mRNA (Perkins et al. 2015). A large number of changes occur in both the brain and female reproductive tract in response to mating and SP that is contained within seminal fluid and subsequently held by the spermatheca (Ito and Tomioka 2016; Wilson et al. 2017; Peng et al. 2005; Avila et al. 2010; Findlay et al. 2014; Ram and Wolfner 2009; Claudia Fricke et al. 2013; C. Fricke et al. 2009; Chapman et al. 2003; Rezával et al. 2012; Yang et al. 2009). Therefore, we also tested whether mating would affect the response to Oa in both WT controls, and flies in which *Oamb* was knocked down using RNAi. *Oamb* knock down significantly blunted the maximum amplitude of the RCaMP signal seen in response to Oa in the spermathecae of mated flies; this effect was not statistically significant for virgin flies (Fig. 5E). Both mated and virgin flies showed a decrease in the duration of the calcium signal seen in response to Oa (Fig. 5F, shown as the time from the onset of the increase until the maximum). We also observed a decrease in the baseline RCaMP signal, although this effect was significant for virgin but not mated flies (Fig. 5G). Together these data indicate that *Oamb* regulates both the calcium level and response to Oa of spermatheca secretory cells although its role may differ between virgin and mated flies.

It is also possible that the function and/or expression of *Oamb* is not uniform across all secretory cells in either mated or virgin flies. While RCaMP appeared to be below the limit of detection in some cells following *Oamb* knock-down (Fig. 5H,I), others appeared to show a clear baseline signal in both mated and virgin flies (Fig. 5H,I), and perhaps a limited response to Oa (Fig. 5I), thus resulting in an inconsistent signal across the circumference of the gland compared to flies expressing WT levels of *Oamb* (Fig. 5D). These inconsistencies likely result from variations in the level of *Oamb* knock down; however, at present we cannot not rule out the possibility that other receptors contribute to the response of the secretory cells to Oa.

### Hyperactivating Oa receptor expressing neurons induces defects in egg laying behavior

We next determined how neurons expressing each Oa receptor might affect reproductive processes required for female fertility. We expressed the temperature-sensitive cation channel, dTrpA1, with the panel of Oa receptor MiMIC Gal4 lines and tested egg laying behavior at 29°C, a temperature at which dTrpA1 activates neurons (Rosenzweig et al. 2005; Hamada et al. 2008). We hypothesized that activation of some if not all of the OaRNs might promote oviposition, similar to previously shown roles for Oa signaling in promoting ovulation (Deady and Sun 2015; White, Chen, and Wolfner 2021; Pang et al. 2022; Cole et al. 2005; Yoshinari et al. 2020). However, we found that activation of OaRNs in female flies led to a significant decrease in egg laying rate compared to controls (Fig. 6, A). We did not detect a change in egg laying rate in control flies expressing any of the Gal4 drivers +/ - *UAS-dTrpA1* and maintained at the permissive 22°C (Sup. Fig. 3).

**Figure 6.**
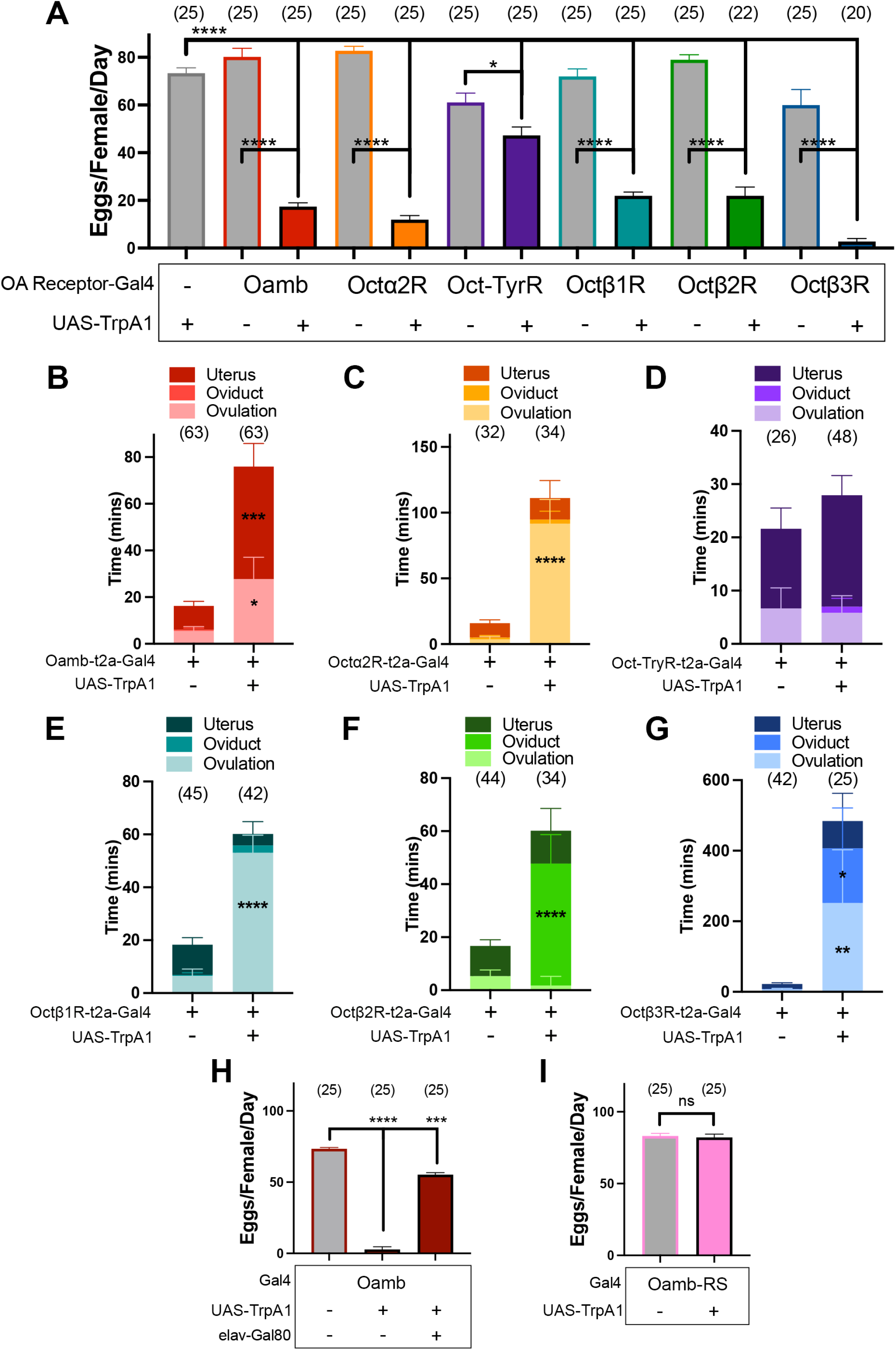
Activating circuits expressing each different octopamine receptor impairs distinct aspects of egg laying. **A.** Expression of *UAS-dTRPA1* via all Oa receptor MiMIC Gal4 drivers during a 48 hour egg laying period significantly decreases counts of eggs laid per female per day **B-G.** Activating neurons expressing each receptor had distinct effects on ovulation and/or passage through the reproductive tract, including retardation of both ovulation and passage through the uterus (**A**, *Oamb*), ovulation alone (**C**, *Octα2R* and **E**, *Octβ1R*), oviduct passage alone (**F,** *Octβ2R*), or ovulation and oviduct passage (**G**, *Octβ3R*). Activation of *Oct-TyrR* cells (**D**) did not show statistically significant changes to any particular step of egg laying despite trending toward retardation of ovulation and oviduct passage. **H.** *Elav-Gal80* co-expression changes the effect of *Oamb* cell hyperactivation on egg laying. **I.** Hyperactivation of oviduct epithelial cells using *Oamb-RS-Gal4* to drive *UAS-dTRPA1* has no effect on egg laying.

Egg laying consists of ovulation (release of eggs from ovary), egg transport through the oviduct, and oviposition (releasing eggs from the uterus to the external substrate). To determine which steps of the egg-laying process might be affected by activating each cell type via *UAS-dTrpA1*, we assayed the distribution of eggs in the female reproductive tract and calculated the average time of each egg spent in ovulation, the oviduct, or in the uterus.

When *Oamb*(+) cells were activated, the time required for both ovulation and oviposition was significantly longer than controls (Fig. 6, B), indicating that these females exhibit a retention of eggs in both the ovaries and uterus. Activating either *Octα2R*(+) or *Octβ1R*(+) cells impaired ovulation but not oviduct passage (Fig. 6, C, E). Conversely, activating *Octβ2R*(+) cells disrupted oviduct passage but did not detectably alter ovulation (Fig. 6, F). Activation of *Octβ3R*(+) neurons increased the time required for both ovulation and oviduct transport (Fig. 6, G). For *Oct-TyrR*(+) cells, we observed a slightly longer average time for the overall egg-laying process with a trend toward increased time spent in the uterus but do not detect a significant change in the time spent in any stage of egg passage (Fig. 6, D). Overall, these findings support the idea that octopaminergic circuits regulate multiple aspects of egg-laying. However, our data now suggest that in addition to *Oamb* and *Octβ2R*, *Octβ1R, Octβ3R, Octα2R* and *Oct-TyrR* expressing cells may also contribute to the regulation of oviposition. They also suggest that Oa signaling may inhibit as well as promote egg laying via some pathways.

*Oamb* expression in multiple non-neuronal cell types in the reproductive tract is required for egg-laying (E. M. Knapp, Deady, and Sun 2018; Lim et al. 2014b; H.-G. Lee, Rohila, and Han 2009; H.-G. Lee et al. 2003; Li et al. 2015). To complement these studies, we aimed to determine whether expression of *UAS-dTrpA1* in neurons is required for the decrease in egg laying we observed following activation of the all *Oamb*(+) cells. To test this, we utilized the temperature sensitive *elav- Gal80* transgene to inhibit Gal4-induced expression of the hyperactivating channel in neurons, and again assayed egg laying. Our results show that *elav-Gal80* suppression of *Oamb-T2A-Gal4* in *UAS- dTrpA1* hyperactivation experiments produces near-control egg laying rates (Fig. 6, H). (Differences between *elav-Gla80* expressing flies and controls may be due to either incomplete inhibition of *dTrpA1* expression in *Oamb*(+) neurons or a contribution by *elav*(-), *Oamb*(+) cells.) We confirmed that expression of *dTrpA1* by the oviduct epithelium driver *OAMB-RS-Gal4* has no effect on egg laying (Fig. 6, I), consistent with the idea that, unlike neurons, non-excitable cells are not responsive to depolarization. These data indicate that neurons as well as non-neuronal cells that express *Oamb* may play a role in regulating oviposition.

### Ovarian egg retention following hyperactivation of octopamine receptor expressing cells

An increase in the time required for ovulation (time within the ovary) might be caused by defects either in follicle cell rupture and/or oocyte development. To differentiate between these possibilities, we examined the content of the ovaries after egg laying. We observed a significant retention of stage 14 mature follicles (Fig. 7, A) following activation of cells expressing *Oamb, Octα2R, Octβ1R, Octβ2R* and *Octβ3R*, but not *Oct-TyrR*. Thus, although Oamb activation in follicle cells promotes ovulation, activation of other octopamine receptor-expressing cells has the opposite effect and appears to impede ovulation. Since *dTrpA1* acts to depolarize cells, the primary candidates for these effects are most likely neurons in the post-mating circuit that provides octopaminergic signals to the ovary. These include processes shown to directly innervate the ovary (Fig. 1) as well as upstream octopaminergic signaling events in the CNS that have the potential to indirectly regulate ovulation. Our data coupled with previous studies using *dTrpA1* suggest that both the direct and indirect effects we observe are due to changes in neuronal signaling; however, we cannot rule out the possibility that other electrically active cells could play a role.

**Figure 7.**
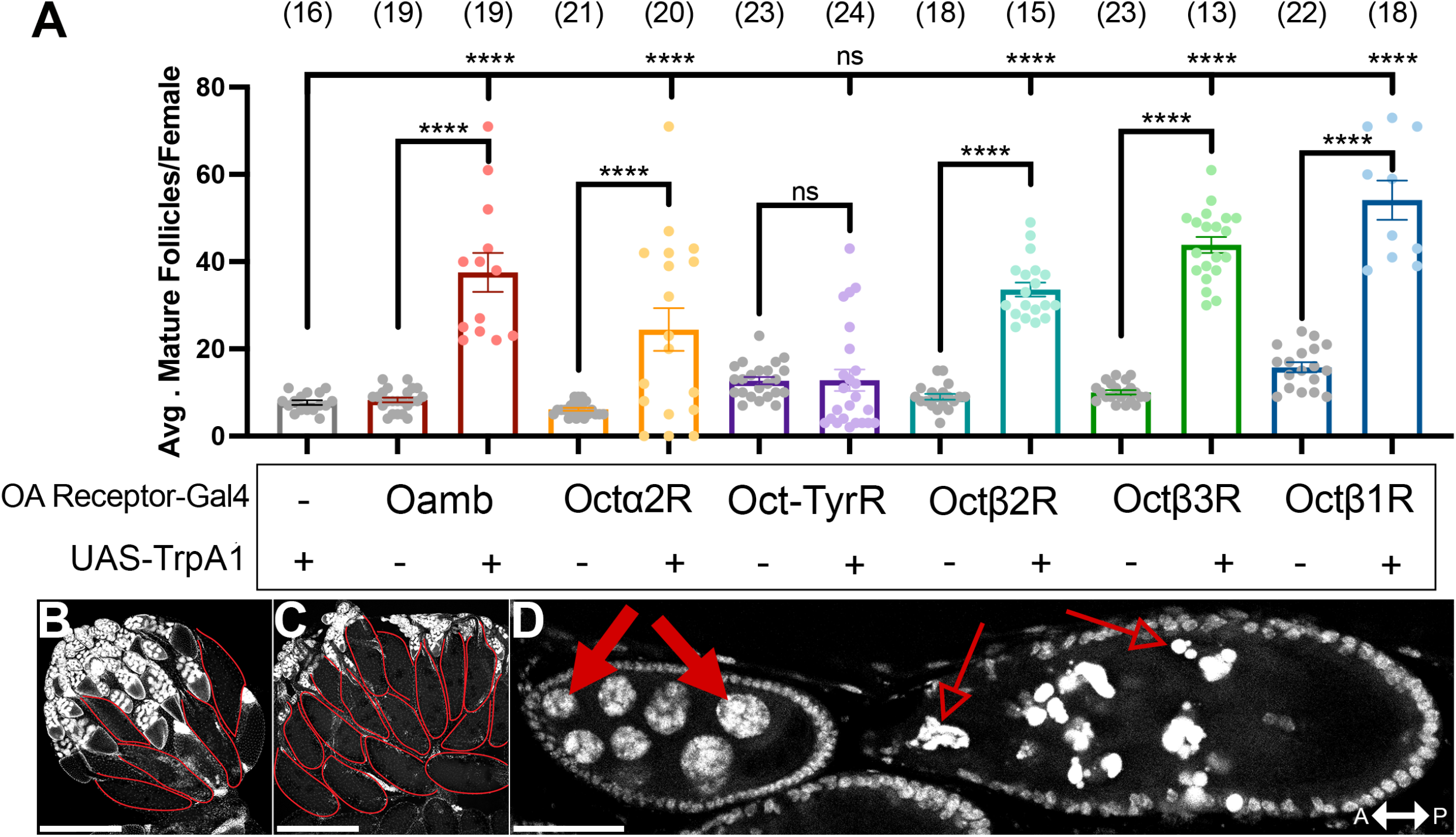
Mature follicles develop but fail to ovulate when octopamine receptor expressing neurons are hyperactivated. **A.** Mature follicle counts per female at time of dissection following a 48hr mating period. **B.** Example ovary from control dataset with 5 mature follicles (red ovals). **C.** Example ovary from dTrpA1-expressing dataset with 16 mature follicles (red ovals). **D.** In some flies in which *Octα2R(+)* and *Oct-TyrR(+)* neurons were hyperactivated, DAPI stains reveal that mid-stage follicles appear to experience nuclear fragmentation in nurse cells. Solid red arrows indicate normal nuclei in an example ovariole. Empty red arrows indicate fragmented nuclei in the same ovariole. Scale bars = 200 μm (**B, C**), 50 μm (**D**).

In sum, our data suggest that some processes regulated by octopamine receptor expressing cells may retard rather than induce ovulation, and that some of these effects are due to neuronal activity. These observations stand in contrast to previous studies in which Oamb activation in non-neuronal cells promoted ovulation via follicle cell rupture and increased the number of germ cells (Deady and Sun 2015; Yoshinari et al. 2020).

### Activation of Octα2R or Oct-TyrR expressing cells induces follicular atresia

In contrast to the Oa receptor expressing cells that we tested, we found that activation of *Oct-TyrR*(+) cells did not lead to a retention in mature follicles (Fig. 7, A). We also observed that activation of *Octα2R* expressing cells elicited mature follicle retention in only about half of the females. Considering both genotypes displayed a decrease in egg laying, we decided to further examine the ovaries to probe for any defects in oogenesis. DAPI staining revealed signs of cell death during mid-oogenesis with both genotypes displaying nurse cell nuclei fragmentation in stages ∼6-9 (Fig. 7, B). Germline cell death in mid-oogenesis can be a result of developmental abnormalities, environmental stress, or drug treatment (Jenkins, Timmons, and McCall 2013). Nutritional deprivation and more specifically protein starvation is the most prominent cause of mid-oogenesis induced cell death (Barth et al. 2011). This starvation induced response slows oogenesis to save costly nutritional resources and is known to be partially regulated by insulin and ecdysone signaling pathways (Terashima et al. 2005; Burn et al. 2015; Pritchett and McCall 2012). Together with these studies, our results suggest that some egg laying effects caused by hyperactivating *Oct-TyrR* or *Octα2R* expressing cells may be due to CNS changes in nutrition or hormonal regulation and that these effects primarily impact follicle development rather than ovulation of mature follicles.

### Hyperactivating glutamatergic/cholinergic but not octopaminergic neurons inhibits egg laying

To explore the mechanism by which Oa receptor expressing cells may regulate egg development and oviposition, we used additional drivers to express *UAS-dTrpA1* in subtypes of neurons that express Oa receptors. As shown in Fig. 3 and (Yoshinari et al. 2020), *ppk*(+) cells and *ChAT*(+) cells in the reproductive tract express multiple subtypes of Oa receptors. We have shown that a subset of glutamatergic neurons that localize to the AbG and project to the reproductive tract also express Oa receptors (Deshpande et al. 2022). To test the possibility that Oa receptors could mediate a reduction in egg-laying via activation of glutamatergic or cholinergic pathways, we expressed *UAS-dTrpA1* in *ppk1.0*(+), *VGluT*(+) or *ChAT*(+) cells. Following activation of cholinergic neurons or *ppk1.0*(+) neurons, we observe a decrease in egg laying rate (Fig. 8, yellow, pink) and an increase in ovulation and oviduct passage time (Fig. 8, D) similar to the activation of cells expressing the receptors *Octα2R* and *Octβ1R*. Activation of *VGluT*(+) glutamatergic neurons resulted in a significant decrease in egg laying rate and an increase in oocyte retention time in the ovary and oviducts similar to the effects of cells expressing *Octβ2R* and *Octβ3R* (Fig. 8A, green, C). By contrast, we failed to detect any decrease or increase in egg-laying when we expressed *UAS-dTrpA1* in octopaminergic neurons using the driver *Tdc2-Gal4* (Cole et al. 2005). Similarly, others have shown that *UAS-dTrpA1* expressed in Tdc2 and Tβh neurons does not elevate egg-laying beyond WT levels, although it rescues the reduction in oviposition caused by exposure to parasitoid wasps (Pang et al. 2022).

**Figure 8.**
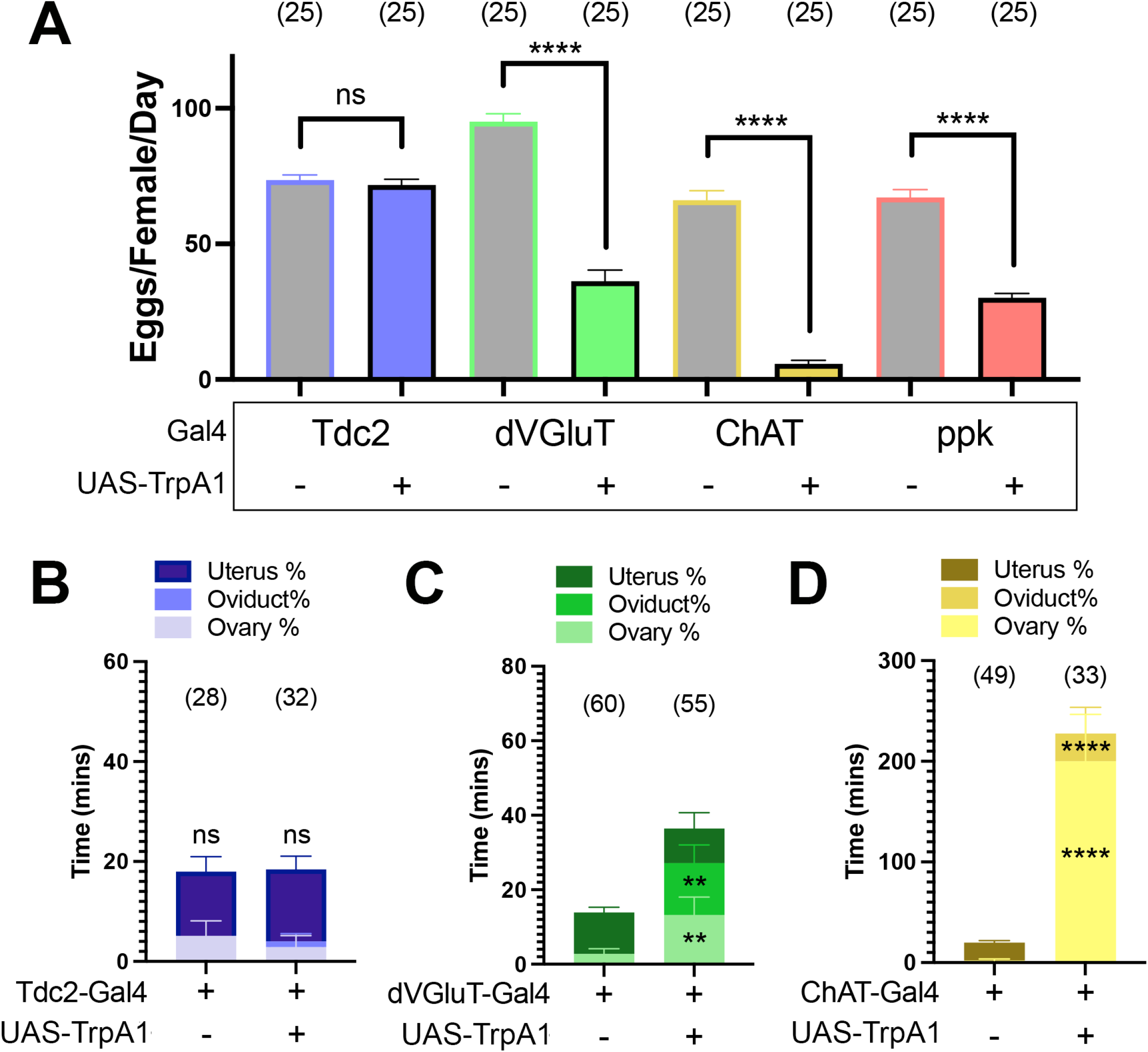
Hyperactivating other cell populations known to partially overlap with octopamine receptor expression produces varied effects on egg laying. **A.** Averages of eggs laid per day over a 48-hr egg laying assay using *UAS-dTrpA1* to hyperactivate cells expressing each indicated Gal4 driver suggest that hyperactivating *Tdc2*(+) cells (blue) has no effect on egg laying and while hyperactivating *dVGluT*(+) (green), *ChAT*(+) (yellow), or *ppk*1.0(+) (red) cells produces egg laying deficits. B. *Tdc2*(+) cell hyperactivation does not significantly alter the amount of time it takes eggs to pass any egg laying process. C. *dVGluT*(+) cell hyperactivation induces defects in both ovulation and oviduct passage. D. *ChAT*(+) cell hyperactivation produces more pronounced ovulation and oviduct passage defects.

### Silencing OaRNs promotes egg laying behavior in both mated and virgin flies

If one of the functions of OaRNs is to inhibit oviposition, then blocking the function of these cells could potentially promote oviposition. To test this hypothesis, we used the Oa receptor MiMIC lines to express the inward rectifying Kir2.1 channel (*UAS-Kir2.1*) and thereby dampen neuronal excitability (Johns et al. 1999). We utilized *tubGal80^ts^* to specifically express *Kir2.1* during egg-laying experiments and avoid the effects of silencing OaRNs during development. When flies are shifted to 29°C to facilitate *Kir2.1* expression, we observed an increase in egg-laying for flies expressing Kir2.1 in *Octα2R*, *Octβ2R*(+), or *Octβ3R*(+) cells (Fig. 9, A, orange, green, blue) but not *Oamb*(+) or *Octβ1R*(+) cells (Fig. 9, A, red, teal). To further test whether inhibiting Oa receptor-expressing cells could promote oviposition, we utilized the transgene *UAS-shibire^ts^* (*UAS-Shi^ts^*), a temperature sensitive mutant form of dynamin, that inhibits neurotransmission via blockade of the exocytotic cycle (Kitamoto 2001). We again observed a slight increase in egg laying rate when cells expressing *Octα2R* or *Octβ2R* were silenced (Fig. 9, B, orange, green respectively), but did not detect any increase following silencing of cells expressing *Octβ1R* or *Octβ3R* (Fig 9, B, teal, blue respectively). When *Oamb-T2A-Gal4/UAS-shibire^ts^* flies were shifted to the restrictive temperature, most females died during the first 24 hrs (data not shown) precluding further analysis. We did not test *Oct-TyrR* in these experiments because hyperactivating *Oct-TyrR* expressing cells had little detectable effect on egg laying and *Oct-TyrR* does not appear to be expressed in *SPR*(+), *ppk1.0*(+) neurons in the uterus (Fig. 6 and data not shown).

**Figure 9.**
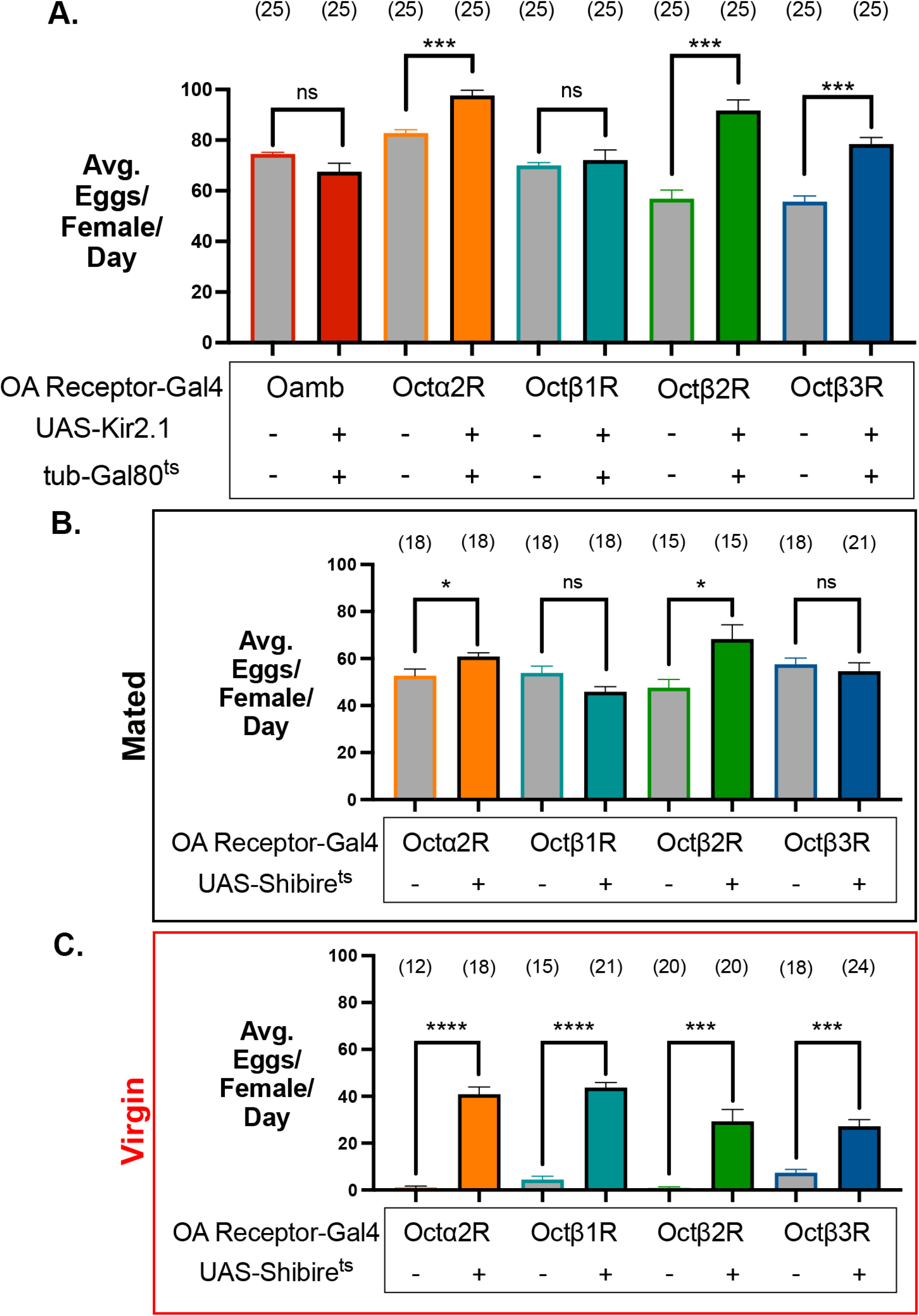
Silencing circuits expressing octopamine receptors increases egg laying in both virgin and mated flies. Oa receptor MiMIC Gal4’s were used to drive *UAS-Kir2.1* (**A**) or *UAS-Shibire^ts^* (**B, C**). At the permissive temperature, egg laying increased in some mated lines (**A**, **B**) and all virgin lines (**C**). Egg laying did not significantly decrease in any condition.

One of the most prominent behavioral changes that occurs after mating is an increase in egg laying (Yang et al. 2009; Rezával et al. 2012; 2014; Yoshinari et al. 2020; Chapman et al. 2003; Ram and Wolfner 2009; Avila et al. 2012; White, Chen, and Wolfner 2021). If one of the functions of OaRNs is to inhibit oviposition, this activity might be expected to be particularly important in virgin females. To test this hypothesis, we repeated our experiment expressing *UAS-Shi^ts^*in OaRNs using virgin rather than mated females. We observe a dramatic increase in egg-laying in all of the cell types we tested including those expressing *Octα2R, Octβ1R, Octβ2R*, and *Octβ3R* (Fig. 9, C). We recognize that daily egg deposition counts can become increasingly variable as egg laying increases (Sup Fig. 3, A). This caveat aside, these data further suggest that in addition to activating processes in the reproductive tract that promote oviposition such as follicle cell rupture, Oa may play an important role in inhibiting oviposition, especially under conditions in which egg-laying would not be productive such as in virgin flies.

## DISCUSSION

It has been known for decades that Oa regulates female fertility and the oviposition circuit in *Drosophila* and other insects (Lange 2009; Tamashiro and Yoshino 2014; White, Chen, and Wolfner 2021; Monastirioti, Charles E. Linn, and White 1996; Zheng et al. 2021; Meiselman, Kingan, and Adams 2018). In *Drosophila*, some loss of function mutants that disrupt Oa synthesis and/or release share a common phenotype marked by retention of mature oocytes in the ovaries (Deady and Sun 2015; Cole et al. 2005; Monastirioti, Charles E. Linn, and White 1996; Monastirioti 2003; Andreatta et al. 2018). Similarly, studies utilizing mutations in *Oamb* and *Octβ2R* have demonstrated a decrease in ovulation and retention of mature follicles within the ovaries (H.-G. Lee et al. 2003; H.-G. Lee, Rohila, and Han 2009; Lim et al. 2014b; Li et al. 2015). *Oamb* also regulates sperm storage, secretory cell activity, oviduct dilation (Avila et al. 2012; Middleton et al. 2006; D. S. Chen, Clark, and Wolfner 2022) and germline stem cell proliferation (Yoshinari et al. 2020; Hoshino and Niwa 2021), while *Octβ2R* is required for lateral oviduct contractions (Deshpande et al. 2022). *Drosophila* also express four other Oa receptors in addition to *Oamb* and *Octβ2R* (Balfanz et al. 2005; Maqueira, Chatwin, and Evans 2005; Qi et al. 2017; McKinney et al. 2020) but their expression patterns and function in the reproductive system have remained speculative.

To gain further insight into the mechanisms by which Oa may regulate oviposition, we have used a panel of high-fidelity Gal4 “MiMIC” lines to map expression in the reproductive tract of all the known *Drosophila* Oa receptors. We have previously shown that *Oamb* and *Octβ2R* are expressed in peripheral neurons proximal to the oviducts and uterus as well as central neurons that project from the AbG to the reproductive tract (Deshpande et al. 2022). Here we show that multiple peripheral neurons that localize to the reproductive tract also express *Octα2R*, *Octβ1R, Octβ3R* and *Oct-TyrR*. These include cell bodies proximal to the oviducts in the MAN and embedded in the musculature of the uterus. Most of these co-express *ChaT* and label with the neuronal marker nervana/anti-HRP. Exceptions include a small subset of cells in the posterior uterus that do not appear to label with anti-HRP; these may represent a novel cell type that is morphologically similar to neurons, but molecularly and/or functionally distinct. We note that some non-neuronal interstitial cells in the mammalian GI and reproductive system share similar properties and play an important role in regulating visceral muscle contractions (Sanders 2019; Dixon et al. 2019). It is possible that insects, and perhaps other invertebrates express an ortholog of this cell type, but this remains unclear.

Using a restricted *ppk1.0-LexA* driver known to express in a subset of afferent *SPR*(+) neurons, we confirm that one anterior uterine cluster of at least three peripheral neurons in the post-mating circuit also co-expresses the cholinergic marker *ChaT* (Yoshinari et al. 2020) and one or more Oa receptors. Of these neurons, one seems to express *Octα2R, Octβ1R*, and *Octβ3R* (Fig. 2, “2”), one seems to express *Octα2R* and *Octβ1R* (Fig. 2, “1”), and one seems to express only *Octβ3R* (Fig. 2, “3”). Other neurons that appear to localize to the same cluster but are *ppk1.0*(-) also express *ChAT* and one or more Oa receptors. These may be part of the broader subset of *ppk1* expressing cells that are not labeled by *ppk1.0-LexA*. The expression of different Oa receptors in different *ppk1.0(+)* neurons raises the possibility that cells in the post mating circuit are sensitive to octopamine. Oa signaling to these cells could represent a negative feedback loop to octopaminergic cells in the AbG that remains active until mating occurs. Expression of differing profiles of Oa receptors further suggests that individual peripheral neurons may respond differently to octopaminergic stimulation, and that the post-mating circuit might be divided into functional units that differ by their expression of different Oa receptor subtypes. Further experiments using intersectional drivers may be useful to test this hypothesis and the function of each cell type. Potential differences between the functions of each OaRN subtype are further highlighted by differences in bouton morphology as well as the degree to which they innervate each organ within the reproductive tract (see Sup. Fig. 1). These differences underscore the possibility that Oa may modulate multiple processes within the reproductive tract beyond its canonical role in promoting oviposition. We suggest that future studies may use Oa receptor expression to help distinguish pathways that regulate female fly fertility.

Live imaging experiments will also be needed to determine how OaRNs respond to Oa, but movement of the reproductive tract following addition of Oa has hampered our attempts to address this question (data not shown). Live imaging of OaRNs in the reproductive tract cells may be possible if the cells are partially or fully dissociated from muscle.

Other cell types proposed to express Oa receptors include follicle cells that surround the developing oocyte and epithelial cells that line the lumen of the oviducts (Deady and Sun 2015; Sun and Spradling 2013; Lim et al. 2014; Lee, Rohila, and Han 2009; Lee et al. 2003; Li et al. 2015; White, Chen, and Wolfner 2021). We also show that cells which line the lumen of the seminal receptacle express *Oamb*, similar to its expression in the epithelium of the oviducts (Deshpande et al. 2022; Lim et al. 2014; Li et al. 2015; Lee, Rohila, and Han 2009; Lee et al. 2003) and that Ca^2+^ levels in the muscle of the seminal receptacle are sensitive to Oa. These results suggest that the epithelial cells of the seminal vesicle may indirectly control the surrounding muscle similar to the mechanism previously proposed for the oviducts (H.-G. Lee et al. 2003; H.-G. Lee, Rohila, and Han 2009; Lim et al. 2014b). We are intrigued by the appearance of wave-like patterns in the seminal receptacle Ca^2+^ activity and speculate that this may play a role in sperm movement within the lumen of the organ, similar to the function of muscle contractions in the movement of eggs within the oviducts. The mechanism(s) which activation of Oamb in the epithelium regulates muscles in either the oviducts or seminal receptacle may be related to indirect pathways for muscle contraction seen in mammals, (H.-G. Lee et al. 2003; H.-G. Lee, Rohila, and Han 2009; Lim et al. 2014b).

### Acute effects in the oviducts and sperm storage organs

Previous results indicate that loss of *Octβ2R* blocks contraction of the lateral oviducts and optogenetic activation of *Octβ2R* expressing neurons can induce lateral oviduct contractions (Deshpande et al. 2022). We find that optogenetic activation of *Octβ1R* and *Oct-TyrR* expressing neurons can also induce lateral oviduct contractions. Since mutation of *Octβ2R* essentially blocks contractions caused by bath applied Oa, the possibility that *Octβ1R*, *Octβ2R*, and *Oct-TyrR* represent three equally important, parallel pathways within the reproductive tract that mediate oviduct contraction seems unlikely. Rather, we speculate that *Octβ1R* and *Oct-TyrR* are more likely to be active in neurons within the CNS and upstream of *Octβ2R*. Alternatively, it remains possible that some of the cells that express *Octβ2R* also express *Octβ1R* and *Oct-TyrR*, but that only the function of *Octβ2R* is required for contractions. Further co-labeling studies and the development of mutations in *Octβ1R* and *Oct-TyrR* will help to distinguish between these possibilities.

While previous studies have demonstrated a requirement for Oa in the regulation of sperm storage, the more acute effects of octopaminergic signaling in sperm storage organs have been less clear. We show that Oa induces calcium transients in secretory cells of the spermathecae and that this effect is blocked by knockdown of *Oamb* within these cells. These data are consistent with a previously assigned role for *Oamb* in sperm storage (D. S. Chen, Clark, and Wolfner 2022; Avila et al. 2012). Based on the results obtained from secretory cell knockdown of *Oamb*, these effects appear to occur via cell autonomous mechanisms within the secretory cells. Similarly, *Oamb* expression is required in the follicle cells for Oa-dependent rupture (Deady and Sun 2015). By contrast, Oa receptors in the epithelium and neurons may regulate muscles in the oviducts via indirect, cell non-autonomous mechanisms (H.-G. Lee et al. 2003; H.-G. Lee, Rohila, and Han 2009; Lim et al. 2014b; Deshpande et al. 2022) (however, see Li et al, 2015), and our current results suggest that *Oamb* may regulate muscles in the seminal receptacle indirectly. These mechanistic differences underscore the complexity of the octopaminergic, regulatory pathways in the oviposition circuit and the potential for divergent outputs.

The relatively high sensitivity of the spermatheca cells to Oa compared to the oviducts (Deshpande et al. 2022), may reflect differences in the relative affinity of Octβ2R versus Oamb, or perhaps differential access of the receptors to bath applied Oa. Concentration-dependent effects have also been observed in the reproductive tracts of other insect species exposed to Oa (Abdoun et al. 1995; Wong and Lange 2014; Lange 2009; Xu et al. 2017).

### Systemic effects in oviposition uncovered using gain of function transgenes

Our functional data using dTrpA1 indicate that activating OaRNs can impede ovulation and egg laying. We confirmed these effects using Kir2.1 and Shi^ts^ to inhibit cells that express Oa receptors and observe an increase in egg-laying. We were initially surprised by these data since previous studies have focused on octopaminergic processes that appear to facilitate ovulation and oviposition (Pang et al. 2022; D. S. Chen, Clark, and Wolfner 2022; White, Chen, and Wolfner 2021; Lim et al. 2014b; Li et al. 2015; H.-G. Lee et al. 2003; Middleton et al. 2006; Monastirioti 2003; Monastirioti, Charles E. Linn, and White 1996; Deady and Sun 2015; Meiselman, Kingan, and Adams 2018). We suggest that our use of gain of function transgenes to activate neurons were important for uncovering effects that are less obvious using loss of function receptor mutants and RNAi transgenes. We also suggest that our use of gain of function strategies was important for probing oviposition because the same gene products may be active in multiple, sequential processes and subject to epistatic effects. In particular, the epistatic relationship between follicle rupture and other processes involved in oviposition may require the use of gain of function methods. For Oa signaling mutants in which oocytes never leave the ovary, downstream effects in the uterus may be difficult or impossible to detect. Therefore, we speculate that the unusual uterine retention phenotype that we report may be absent in loss of function *Oamb* mutants because the oocytes are trapped at an upstream site in the ovary (Deady and Sun 2015; H.-G. Lee et al. 2003; H.-G. Lee, Rohila, and Han 2009; Yoshinari et al. 2020). By contrast, the use of gain function probes to activate downstream processes allowed us to detect the uterine retention phenotype. Similarly, oviduct retention might be occluded by upstream retention of mature follicles in the ovaries. If so, retention of eggs in the ovary seen with *Octβ2R* knock-down or mutants might have occluded downstream effects in the oviducts (Lim et al. 2014b; Li et al. 2015).

Although the epistatic relationship is less obvious, we suggest that some of the effects we observe may reflect disruption of processes upstream of follicle rupture. Instances of follicular atresia observed following hyperactivation of *Octα2R* and *Oct-TyrR* expressing neurons suggest that the underlying circuits regulate follicle development. Based on the phenotype reported in other genetic studies, we speculate that the follicular atresia we observe could be caused by disruption of nutritional intake or other upstream homeostatic circuits (Jenkins, Timmons, and McCall 2013; Barth et al. 2011; Terashima et al. 2005; Burn et al. 2015; Pritchett and McCall 2012). Though this phenotype seems to be only partially penetrant, it might occlude some of the effects of *Octα2R* and/or *Oct-TyrR* on downstream egg laying processes. The lack of any obvious impairment to follicular development in hyperactivation assays involving the other Oa receptors, however, indicates that the effects we observe for most Oa receptor expressing cells are likely due to direct disruption of the reproductive tract rather than broader, metabolic mechanisms that impact multiple organ systems and influence reproductive activity indirectly.

### Possible cell and molecular mechanisms

CNS circuits that regulate oviposition include pCL1 neurons in the brain that innervate oviposition descending neurons (oviDNs) (F. Wang et al. 2020; Feng et al. 2014). It is tempting to speculate that OaRNs might regulate pCL1 or oviDN, or additional excitatory or inhibitory neurons within the same circuit (F. Wang et al. 2020). We are particularly drawn to the observation that hyperactivation of *Oamb*(+) neurons results in retention of eggs in the uterus just prior to deposition (Fig. 6, B). Following follicle rupture, eggs ovulate and pass through the oviduct in a very short amount of time in WT flies and without significant delay under baseline conditions (Mattei et al. 2015). By contrast, flies can retain fertilized eggs in their uterus until sensory inputs indicate that egg laying can occur in a predator/toxin free environment (Pang et al. 2022). We speculate that *Oamb*(+) neurons in the CNS may represent a behavioral choice point and help to regulate the decision to deposit eggs. Further studies of the CNS connectome combined with single cell sequencing, and optogenetics will be needed to test this hypothesis and identify the underlying circuits.

It remains unclear why hyperactivation of presynaptic *Tdc2*(+) neurons with dTrpA1 does not appear to increase egg-laying. Similarly, *dTrpA1* expression in *Tdc2*(+) and *Tβh*(+) neurons rescued a reduction in oviposition caused by exposure to parasitoid wasps, but did not elevate egg-laying beyond WT levels (Pang et al. 2022). We speculate that, if the effects of Oa are as complex as we suggest and it acts to both promote and retard ovulation and oviposition, simultaneous activation of all octopaminergic pathways might not appear to have any effects under some conditions. It is also possible that, under some circumstances, octopamine and tyramine have opposing effects in oviposition as they do for larval locomotion (Saraswati et al. 2004). Further experiments using intersectional drivers that are specific for subsets of octopaminergic neurons may be needed to detect a net loss or gain in fertility in the absence of exogenous stimuli such as threats from parasitoid wasps (Pang et al. 2022). A previously described intersectional approach using *doublesex* is useful for expression in the multicellular cluster of octopaminergic neurons that innervates the reproductive tract, but cannot be used to stimulate individual octopaminergic neurons within the cluster (Rezával et al. 2014).

Further experiments also will be needed to determine which post-synaptic neurons that express specific Oa receptors are responsible for the effects we observe. We are intrigued that the effects of hyperactivating or silencing OaRNs on egg laying are similar to those seen in experiments involving the *SPR*(+) neurons of the post-mating circuit (Yoshinari et al. 2020; Yang et al. 2009; Yapici et al. 2008). We speculate that some of neurons in the peripheral *SPR*(+) post mating circuit that express *ppk1.0* and Oa receptors could be responsible for the inhibition of egg laying we observe. Since SPR is coupled Gα_i_ or Gα_o_ and nominally inhibitory (Yapici et al. 2008), opposing Oa inputs mediated by Oa receptors coupled to Gα_s_ or Gα_q_ could potentially activate one or more subsets of neurons in the post-mating circuit until SP silences their activity. This remains highly speculative, but further intersectional studies using SPR and OaRN drivers might be used to probe for more specific subsets of *SPR*(+) neurons and to test these hypotheses.

Importantly, all the transgenes we have used here act by directly activating or inhibiting neuronal activity rather that activating or inactivating Oa receptors. Oa receptors, like most other GPCRs can have net “inhibitory” or “excitatory” effects which can vary across cell types, downstream effectors and the subcellular location of the receptors (Robb et al. 1994; M. Wang et al. 2007). The coupling of Oa receptors to nominally “excitatory” G proteins (Gα_s_ and Gα_q_) and their downstream effectors has been extensively examined *in vitro* and in the epithelial cells within the reproductive tract (Y.-C. Kim et al. 2013; H.-G. Lee, Rohila, and Han 2009; Debnath, Williams, and Bamber 2022; Xu et al. 2017). We therefore speculate that effects seen in neuron hyperactivation experiments using Oa-receptor drivers may be similar to the effects of increased Oa signaling to these cells. However, further experiments will be needed to more precisely determine the *in vivo* effects of Oa receptors in neurons within the oviposition circuit and CNS as well as how each may influence egg-laying.

## Acknowledgments

We thank Shivan Bonanno, a UCLA postdoctoral scholar in the Krantz lab, for his consultation and support. We thank the labs of Dr. Lawrence Zipursky and Dr. Kelsey Martin at UCLA for graciously sharing their confocal microscopy equipment. We thank Hugo J. Bellen Pei-Tseng Lee and other members of the Bellen lab for generating the MiMIC lines and the GAL4 conversions, especially Yuchun He, Wen-Wen Lin, and Drs. Sonal Nagarkar-Jaiswal and Oguz Kanca as part of their efforts to generate “A comprehensive resource for manipulating the *Drosophila* genome with swappable insertion cassettes”, funded by NIH R24 OD031447, to Hugo J. Bellen and Oguz Kanca. We thank the following people for generously supplying fly lines: Dr. Kyung-An Han (University of Texas, El Paso), Dr. Bing Ye (University of Michigan) and Dr. Robert Kittel (University of Würzberg).

## Data Availability Statement

Fly strains are available upon request and/or from the Bloomington Stock Center as indicated. The authors affirm that all data necessary for confirming the conclusions of the article are present within the article, figures, and tables.

## Figure Legend

**Supplemental Figure 1. Octopamine receptors are expressed by neurons with terminals in reproductive organs.** *Oamb-T2A-gal4* (**A**), *Octα2-T2A -Gal4* (**B**), *Oct-TyrR-T2A-Gal4* (**C**), *Octβ1R-T2A-Gal4* (**D**), *Octβ2R-T2A-Gal4* (**E**), and *Octβ3R-T2A-Gal4* (**F**) were used to drive *UAS-mCD8-GFP* expression. Reproductive systems were dissected and stained with anti-GFP antibody (green) and phalloidin f-actin stain (magenta). Representative nerve terminals were captured in the ovaries (**A-F, i**), calyx regions (**A-F, ii**), common oviduct (**A-F, iii**), seminal receptacle (**A-F, iv**), and uterus (**A-F, v**). The approximate density of innervation in each region by neurons expressing each different receptor type was then quantified by calculating the area innervating nerves took up as a percent of equivalent areas of tissue across different preparations (**G**). From the same images, the approximate size of representative terminals was also calculated (**F**). Scale bars = 100 μm (**A-F, i**) and 10 μm (**A-F, ii-v**)

**Supplemental Figure 2. Octopamine receptors are expressed by cells in the reproductive system that can be labelled with a neuron-specific antibody.** MiMIC Oa Receptor Gal4’s were used to express *UAS-mCD8-GFP*. Signal from GFP (**Aii**, **Bii**, **Cii**) and anti-HRP/nervana (**Aiii**, **Biii**, **Ciii**) antibody labels were compared (**Ai**, **Bi**, **Ci**) in reproductive systems and attached MANs. Representative regions of co-labeling are indicated by chevrons. Scale bars = 200 μM.

**Supplemental Figure 3. Oa-Receptor-T2a-Gal4, UAS-dTrpA1 flies kept at a restrictive temperature of 22°C did not show defects in fertility compared to controls.** Neither egg laying (**A**), egg passage through any specific reproductive organ (**B**), nor mature follicle development (**C**) was significantly reduced in flies harboring the same alleles as in Figure 3 but kept at a temperature restricting TRPA1^ts^ expression. Flies harboring *Octβ3R-t2a-Gal4* and *UAS-TRPA1^ts^* displayed a slight increase in egg laying (**A**, blue). However, their egg laying rate was not significantly different from the Gal4 (-) control (**A**, grey).

**Supplementary Video 1. Transient calcium wave activity in the muscle of the seminal receptacle.** RCaMP1b fluorescence in the muscle of the seminal receptacle presents as waves of increasing signal spontaneously before the addition and after addition of Oa. A global increase in the amplitude of the RCaMP1b signal response is seen after addition of Oa (1μM) at 5 sec. We did not detect a change in the rate of Ca^2+^ transients. The clip has been sped up 3x (12fps to 36fps) and trimmed to show only the 10 sec period bracketing the addition of Oa at 5 sec.

